# *putzig* safeguards genome integrity by contributing to the Piwi-mediated repression of transposon activity in the female germline of *Drosophila* in a two-tiered fashion

**DOI:** 10.64898/2026.03.11.711025

**Authors:** Lea Hüttinger, Ludmilla Kober, Mirjam Zimmermann, Sebastian Schlensog, Marc Engelhart, Anja C. Nagel

## Abstract

Genome integrity in the germline is jeopardized by the activity of transposable elements (TEs). Transposon activity is kept in check by the Piwi-piRNA pathway, which silences TEs post-transcriptionally in the cytoplasm as well as co-transcriptionally in the nucleus. piRNAs derive from long precursor transcripts originating at piRNA-loci by non-conventional transcription. They are processed in the cytoplasm and loaded into Piwi-piRNA complexes that then enter the nucleus to bind to target RNAs and direct local heterochromatin formation at TE loci, involving several downstream effectors.

In this work, we have analyzed the role of Putzig (Pzg) in Piwi-mediated TE silencing in the female germline. Pzg has multi-facetted roles during *Drosophila* development and is important for DNA integrity, germ cell differentiation and survival. Here, we provide evidence for a two-tier activity of Pzg by serving as a hub for various Piwi-pathway members. Firstly, Pzg assists transcription initiation at piRNA clusters by coupling the Trf2-Moonshiner transcription initiation complex to the Rhino-Deadlock-Cutoff complex, thereby promoting formation of piRNA-precursor transcripts. Secondly, in the context of co-transcriptional gene silencing, Pzg licences heterochromatin formation by linking the Piwi-piRNA silencing machinery and the histone demethylase Lsd1, involved in promoter inactivation.

## Introduction

The maintenance of stability and integrity of eukaryotic genomes is key to organismal survival, successful reproduction as well as proper development. Numerous challenges afflict genomic integrity, resulting in genotoxic stress by accumulation of mutations and DNA damage, thus changing critical gene expression programs. Whereas genome damage in somatic cells may induce cancer and hence threatens the health of an individual, respective damage in the germline threatens subsequent generations with long-lasting implications. The ability of transposable elements (TEs) to move within a genome makes them a bona fide intrinsic source to cause detrimental effects on gene and genomic integrity (Bourque *et al*., 2018). TEs are highly abundant mobile genetic elements, which constitute about 45% of the human- and about 20% of the *Drosophila* genome, respectively (Bourque *et al*., 2018; Mérel *et al*., 2020; Kaminker *et al*., 2002). In contrast to humans, where most TEs are considered inactive, 30% of the mobile elements in *Drosophila* are believed to be active. Thus, studying the dynamics and regulation of TE mobilization in *Drosophila* is quite promising to gaining deeper insights into the molecular mechanisms that keep their activity in check thereby defending genome integrity (Kaminker *et al*., 2002; Barrón *et al*., 2014). In the animal germline, the existence of an effective defense system is well established and relies to the Piwi-interacting RNA (piRNA) silencing system, which efficiently silences TEs either via post-transcriptional gene silencing (PTGS) in the cytoplasm or co-transcriptional gene silencing (TGS) in the nucleus (reviewed in Czech *et al*., 2018; Di Stefano, 2022; Claro-Linares and Rojas-Ríos, 2025). The majority of piRNAs is derived from specific genomic loci, called piRNA clusters, harboring inactive transposon insertions, thereby functioning as archives of transposon sequences (Brennecke *et al*., 2007; Senti and Brennecke, 2010). In contrast to the somatic piRNA clusters, germline specific piRNA clusters are dual-stranded, i.e. are typically processed from longer precursors transcribed in a non-canonical manner from both genomic strands (Brennecke *et al*., 2007; Klattenhoff *et al*., 2009; Mohn *et al*., 2014; Andersen *et al*., 2017; reviewed in Senti and Brennecke, 2010; Czech *et al*., 2018; Ozata *et al*., 2019). These dual-strand piRNA clusters exhibit heterochromatic transcriptional silencing signatures such as histone H3 lysine 9 tri-methylation (H3K9me3), however bypass transcriptional silencing by recruiting specific multiprotein complexes that allow transcriptional initiation in a promoter-independent manner even in heterochromatic surroundings. Here, the germline-specific HP1a homologue Rhino (Rhi) is associated with H3K9me3 marks, enabling the assembly of the RDC complex together with Deadlock (Del) and Cutoff (Cuff) (Klattenhoff *et al*., 2009; Mohn *et al*., 2014; Zhang *et al*., 2014; Yu *et al.,* 2018). RDC then allows the recruitment of an alternative basal transcription factor complex consisting of the TFIIA-L paralogue Moonshiner (Moon), TFIIA-S and the TATA box binding protein-related factor 2 (Trf2), which initiates the non-canonical transcription from these heterochromatic regions (Andersen *et al*., 2017).

In *Drosophila* germline cells, the long piRNA precursor transcripts are exported to cytoplasmic perinuclear foci, called nuage, where eventually the small piRNAs associated with members of the Piwi protein family are generated (ElMaghraby *et al*., 2019; Kneuss *et al*., 2019; reviewed in Czech *et al.,* 2018; Ozata *et al*., 2019). In the nuage, the Piwi family members Aubergine (Aub) and Argonaute (Ago) plus several cofactors amplify piRNAs in a so-called ping-pong cycle, by repeated reciprocal cleavage of transposon mRNA and piRNA cluster transcripts (reviewed in Czech *et al.,* 2018; Claro-Linares and Rojas-Rios, 2025). Phased piRNA production, maturation and loading into the Piwi-piRNA complexes occurs on mitochondria by help of Armitage (Armi), Zucchini (Zuc) and further cofactors (Ge *et al.,* 2019; Yamashiro *et al.,* 2020; reviewed in Claro-Linares and Rojas-Rios, 2025). Once loaded with piRNA, Piwi enters the nucleus, where it associates with its target mRNA, i.e. nascent TE transcripts to eventually induce co-transcriptional gene silencing of TEs (Sienski *et al*., 2012; Wang and Elgin, 2011; Le Thomas *et al*., 2013). Target engagement triggers the formation of the activated Piwi complex (Piwi*) together with Maelstrom (Mael) and Asterix (Arx, also called Gtsf1), serving as a hub for downstream silencing effectors (Portell-Montserrat *et al.,* 2025). Via the binding of Nuclear export factor 2 (Nxf2), Piwi* recruits the so-called SFiNX complex including NTF2-related export protein 1 (Nxt1) and Panoramix (Panx). The Panx-Nxf2-Nxt1 co-transcriptional silencing complex bears several other names apart from SFiNX, abbreviated Pandas, PICTS or PPNP (Batki *et al.,* 2019; Zhao *et al.,* 2019; Fabry et al., 2019; Murano *et al*., 2019; reviewed in Di Stefano, 2022). SFiNX functions upstream of the chromatin modifying machinery, whereby Panx initiates co-transcriptional TE silencing presumably by facilitating transcription termination (Sienski *et al*., 2015; Batki *et al.,* 2019; Fabry *et al.,* 2019; Murano *et al.,* 2019; Zhao *et al.,* 2019; Andreev *et al*., 2022; Liu *et al*., 2025). Continuous TE silencing, however, involves chromatin remodellers, notably Eggless (Egg, or SetDB1) and Lysine-specific demethylase 1 (Lsd1, also called Su(var)3-3) that both play important roles in germ cell development and TE silencing (Di Stefano *et al*., 2007; Yoon *et al.,* 2008; Eliazer *et al.,* 2011; Wang et al., 2011; Clough *et al*., 2014; Sienski *et al.,* 2015; Lepesant *et al*., 2020). Whilst the H3K4me1/2 demethylase Lsd1 impedes transcription initiation, Egg-mediated repressive histone methylation marks (H3K9me3) allow the recruitment of Heterochromatin Protein 1a (HP1a), eventually causing persistent chromatin compaction (reviewed in Meyer-Nava et al., 2020; Di Stefano, 2022). The exact mechanisms regarding the recruitment of these chromatin modifiers are incompletely understood, yet, SUMOylation plays an important role for the assembly of the silencing complex, allowing the engagement of Egg/SetDB1 and subsequent H3K9 tri-methylation followed by HP1a-mediated heterochromatin formation (Ninova *et al*., 2020; Andreev *et al*., 2022; Bence *et al.,* 2024). Recruitment of Lsd1 is less well understood, involving Su(var)2-1 (also named *ovaries absent, ova*), which directs global histone deacetylation and couples Lsd1 to HP1a (Yang, *et al*., 2019; Walther *et al.,* 2020). Together, these repressive marks guide effector proteins such as HP1a to TE loci, initiating heterochromatin formation and finally transcriptional silencing of TEs (Le Thomas *et al*., 2013; Iwasaki *et al*., 2016; Fabry *et al*., 2019; Ninova *et al*., 2020; reviewed in Meyer-Nava et al., 2020; Di Stefano, 2022).

In this study, we examined the role of the gene *putzig* (*pzg*, also named *Z4*) in sustaining genome stability in the female germline. The *pzg* gene encodes a zinc-finger containing multifunctional protein. Originally identified as a component of the TRF2/Dref complex in the transcription of replication related genes, it was later shown to be also an important regulator of the activity of several signaling pathways, including Notch, JAK/STAT and EcR, during the development of imaginal tissues (Hochheimer *et al.,* 2002; Kugler and Nagel, 2007; Kugler *et al*., 2011). Moreover, Pzg protein was found in the NURF chromatin remodeling complex and was also involved in chromatin activation and chromatin structure, for example at the telomeres (Eggert *et al*., 2004; Andreyeva, *et al*., 2005; Kugler and Nagel, 2010; Kugler *et al.,* 2011; Silva-Sousa *et al*., 2012; Puerto *et al.,* 2024). Pzg activity is essential for embryonic to larval development and growth. Hence, its role in the germline was assessed by short hairpin RNA interference (shRNAi), demonstrating Pzg’s requirement for germ cell development and female fertility (Kober *et al*., 2019). Downregulation of Pzg not only impaired the differentiation of germline stem cells but also induced cell death in the germarium, thereby generating a new niche like environment by the activation of Dpp and Wnt signals (Kober *et al*., 2019). The primary cause of germline cell death, however, remained unresolved and prompted us to investigate the role of Pzg in transposon silencing. Our results indicate that loss of *pzg* in the germline de-represses transposon activity and thus challenges genome- and DNA-integrity. We provide molecular evidence that Pzg acts in a two-tiered fashion: Firstly, by association with the RDC-Trf2 complex, Pzg stimulates the transcription of piRNA-precursor transcripts. Secondly, in a complex with Piwi and SFiNX, it promotes the recruitment of Lsd1, thereby fostering epigenetic silencing of TE loci. Taken together, we propose that Pzg is an important factor in ensuring genome stability in the germline by combatting transposon activity at multiple levels.

## Materials and Methods

### Key resources are listed in Key Resources Table

#### Fly work

Maintenance of flies was at 18°C; experiments were performed at either 25°C or 29°C. To avoid overcrowding, crosses were set up with 20 females and 15 males each and changed every other day on enriched food as described before (Kober *et al*., 2019). Ovaries from a minimum of 10-15 females were sampled for ovariole analyses by fluorescence - or confocal microscopy. Experiments were repeated at least two times. Fly stocks were recombined and combined according to standard genetics for rescue experiments and reporter analyses. Genotypes were verified by single-fly PCR using specific primer sets. Fly strains and primers are listed in Key Resources Table.

#### Antibody, DAPI or EdU staining of adult ovaries

Ovaries were dissected from 3-5 days old females of the corresponding genotype, and collected in PBS (130 mM NaCl, 7 mM Na_2_HPO_4_, 3 mM NaH_2_PO_4_) on ice.

After 18-20 minutes fixation in 4% paraformaldehyde (PFA), the ovaries were washed at least three times for 15 min with PBX (PBS + 0.3% Triton^®^ X-100). ***For antibody staining***, ovaries were first subjected to a 30-60 min blocking step in PBX and 4% normal goat serum (NGS) at room temperature (RT) followed by antibody incubation over night at 4°C. After several washes with PBX, followed by pre-incubation with 4% normal goat serum, secondary antibodies were added for incubation overnight at 4°C. Finally, after repeated washing steps with PBX, ovaries were mounted in Vectashield^®^ and analyzed either with a Bio-Rad MRC1024 confocal system coupled to a Zeiss Axiophot microscope using LaserSharp 2000 imaging software, or with a Spinning Disk microscope and 3i SlideBook6 software for acquisition. ***For DAPI staining***, the ovaries were stained with 4’,6-Diamidine-2-phenylindol (DAPI) (1 μg/μl in 0.3% PBX) in the dark for 4 min, after the fixation and washing steps described above. Ovaries were then washed again three times for 5 min in 0.3% PBX and mounted in 80% glycerol for documentation on a Zeiss Axioskop 2 plus, connected to a Canon EOS 700 camera. ***For EdU staining***, ovaries were transferred to 1 ml M3/EdU solution (M3 medium, 50 µM EdU) and incubated at RT for 2 hours. After 15 min fixation in 4% PFA, the ovaries were washed five times in PBX. Antibody staining was performed as described above. After incubation with secondary antibodies, the ovaries were washed four times for 10 min in PBX and twice for 10 min in PBS. After dissection, ovaries were mounted in Vectashield^®^ and documented with a Bio-Rad MRC confocal system as described above.

Images were processed and assembled into figures using Fiji (ImageJ2), Corel-Photo Paint and CorelDRAW software.

### Fluorescent in situ hybridization (FISH) with piRNA probes 38C2 and/or 42AB

#### 1. Performing FISH

Fluorescence in situ hybridization (FISH) was used to examine the expression pattern of piRNA clusters *42AB* and *38C2* in *pzg*-deficient nurse cells. For this purpose, crosses between *matα*-Gal4 virgins and UAS-shRNA*-w* and UAS-shRNA-*pzg* males were set up, incubated at 29 °C, and regularly transferred. The *matα*-Gal4 driver induces RNAi expression only in later egg stages, allowing further development of the *matα*-Gal4/UAS-shRNA-*pzg* ovaries. For each approach, 5-10 ovaries of 3-5 days old F1 generation were prepared and gently shaken in fixative solution (4 % PFA, 0.15 % Triton X-100 in PBS) at RT for 20 min. In the following steps, RNase-free working conditions were maintained. The fixative solution was replaced by three ten-minute washing steps with 0.3% PBX. Permeabilization was carried out overnight to a maximum of one week in 70% EtOH at 4°C. The ovaries were then incubated for 5 min at RT in RNA FISH wash buffer [10 % (v/w) formamide in 2x SSC (0.3 M NaCl, 0.03 M sodium citrate in DEPC-H_2_O, pH 7.0)]. Hybridization was performed in 50 µl hybridization buffer (10 % (v/w) dextran sulphate, 10 % (v/w) formamide in 2x SSC) with 0.5 µl RNA FISH probe each, gently rotating overnight at 37 °C in the dark in a hybridization oven. The following day, the ovaries were washed twice for 2 min at RT in the dark in RNA FISH wash buffer. This was followed by staining with 1 µg/µl DAPI in 2x SSC for 10 min at RT. The ovaries were then washed twice for 10 min at RT in 2x SSC, isolated, and mounted in Vectashield®.

#### 2. Evaluation of FISH

Two biological replicates were evaluated per genotype and FISH probe. For this purpose, images of at least two egg chambers in stage 8 were first taken by confocal microscopy (Zeiss LSM 700). The FISH spots were then assigned to the regions: “nucleus”, “nuage” and “cytoplasm” in ImageJ2 (2.16.0) and counted, using the same color balance and contrast settings throughout. A total of at least 55 nuclei were counted per genotype and FISH probe. The box plots were created using the R packages ggplot2 (3.5.2), tidyverse (2.0.0), and dplyr (1.1.4), and significance was determined using the Mann-Whitney-Wilcoxon test.

### Transposon-lacZ reporter assays

3-5 days old female flies were anesthetized with diethyl ether for approx. 5 min, and the ovaries were prepared and collected in PBS (130 mM NaCl, 7 mM Na_2_HPO_4_, 3mM NaH_2_PO_4_) on ice, fixed in 0.5% glutaraldehyde in PBS for 15 min and then washed three times for 15 min in PBS. The ovaries were incubated in a freshly prepared staining solution (1 M MgCl_2_, 5 mM NaCl, 310 mM K_3_[FeIII(CN)_6_], 310 mM K_4_[FeIII(CN)_6_], 10% Triton X100, 8% X-Gal in PBS) for several hours or overnight at 37°C to detect ß-galactosidase activity. The reaction was stopped by washing in 500 µl PBX. The ovaries were then prepared in PBS and mounted in 80% glycerol. Ovaries were documented at the Axiophot stereomicroscope coupled with a Pixera PVC 100C camera and iWorks 2.0 software.

### Feeding of Lamivudine (3TC)

Oral treatment with the nucleoside analogue and reverse transcriptase inhibitor Lamivudine (3TC) was modified based on the protocol by Lin *et al*. (2020). For a final concentration of 10 mM 3TC, 11.5 mg 3TC was mixed with 1.7 g dry yeast and 3.3 ml ddH_2_O. As a control, the dry yeast was mixed with water without 3TC. Yeast paste was struck out in plastic vials containing normal food and 50 embryos each were collected from a clutch of either *nos*-Gal4::UAS-shRNA-*w* (control) or *nos*-Gal4::UAS-shRNA-*pzg* crosses on apple juice plates and directly transferred to the yeast paste. This ensured that development took place exclusively on the prepared yeast paste from the very beginning of food intake up to the preparation of female ovaries.

### Nuclear extracts from L1 larvae

200 mg larvae at first larval instar (L1) were flash frozen in liquid nitrogen and stored at −80°C. Nuclear extracts were prepared according to Maier *et al*. (2002). All steps were performed in the cold room (4°C) and on ice, and all buffers contained Protease inhibitor cocktail. Larvae were homogenized in 800 µl HB-buffer (10 mM HEPES pH 7.6, 10 mM KCl, 1 mM MgCl_2_, 0.5 mM DTT) and filtered through a wet Miracloth filter. In a first step, the dirt was pelleted for 5 min at 200 x g. The supernatant was then submitted to centrifugation for 10 min at 3.000 x g, resulting in a crude nuclear pellet. The pellet was washed twice in 400 µl HB-buffer, sedimented in between as above. Then the pellet was resuspended in 500 µl PA-buffer (20 mM HEPES pH 7.6, 10 mM KCl, 2 mM MgCl_2_, 0.5 mM DTT with 0.3 M sucrose) and centrifuged through a cushion of 500 µl PA-buffer with 1.7 M sucrose for 30 min at 50.000 x g. The nuclear pellet was resuspended in 200 µl WB-buffer [10 mM HEPES pH 7.6, 1 mM MgCl_2_, 1 mM CaCl_2_, 150 mM NaCl] and frozen at −80°C until use. Nuclear extracts were separated on a large SDS-PAGE (17%) for 30 h at 4°C and the relevant gel slice (7-28kDA) was cut out and blotted. The blot was probed with ⍺-H2A and ⍺-γH2AX (pSer139) antibodies.

### Yeast two-hybrid interaction assays

The yeast two-hybrid experiments were based on the Golemis-Brent hybrid system, using EGY48 yeast cells and performed as previously described according to standard protocols (Gyuris *et al*., 1993; Golemis and Brent, 1997; Gahr *et al*., 2019). All interaction experiments were repeated at least three times. cDNA prepared for qRT-PCR (see below) was used for PCR-amplification of potential Pzg interaction partners, cloned in pEG202 vector to produce fusion proteins with the LexA DNA-binding domain, whereas Pzg full length as well the Pzg-subdivision constructs were cloned in pJG4-5 vector possessing the B42 activator domain. Primers used for the amplification and cloning are listed in Key Resources Table. Constructs were sequence verified (Macrogen Europe, Amsterdam, Netherlands). Correct expression of LexA-fusion proteins (pEG-constructs) and HA-fusion proteins (pJG-constructs) was tested by western blot using ⍺-LexA and ⍺-HA antibodies, respectively. pSH18-34 was used as reporter, expressing lacZ upon interaction of tested constructs.

### Protein immunoprecipitation

Protein immunoprecipitation (IP) was used to identify potential interaction partners of Pzg in ovaries. For this purpose, a transgenic fly strain with GFP-labelled Pzg protein (Sarov *et al*., 2016) was used to precipitate Pzg using anti-GFP nanobodies (GFP-Trap^®^) followed by mass spectrometry (IP-MS). In addition, transgenic fly strains expressing tagged candidate genes were used to perform IP’s followed by Western blot analyses to verify possible co-IP with Pzg using antibodies detecting either Pzg or the respective tag.

*Generation of protein extracts from ovaries:* Either 200 (IP-MS) or 400 ovaries (for IP followed by Western Blotting) were prepared in PBS. The ovaries were then homogenized in 450 µl protein lysis buffer [20 mM TRIS-HCl pH 7.5, 150 mM NaCl, 2 mM MgCl_2_, 10 % glycerol, 1 mM DTT, 1 mM Pefabloc® SC-Protease Inhibitor, 0.2 % NP40 in ddH_2_O] in a glass homogenizer with a size B pestle. After centrifugation for 15 minutes at 4°C with 14.000 rpm, the lysate was transferred to a fresh Eppendorf tube and stored at −80°C until further use.

*Performing the IP:* All steps were performed in the cold. Precipitation was carried out using magnetic GFP-Trap^®^/RFP-Trap^®^ or Myc-Trap^®^ agarose beads depending on the genotype of the transgenic fly strains, using a Magnetic Separation Rack. Binding Control agarose beads were used as a negative control. For preparation, 25 µl of the agarose bead mixture was pipetted into low retention Eppendorf tubes and washed three times with washing solution 1 [150 mM NaCl, 50 mM TRIS-HCl pH 7.5, 0.1 % SDS, 1 tablet of protease inhibitor cocktail in ddH_2_O]. For precipitation, 200 µl of protein extract and 300 µl of washing solution 1 were added to the agarose beads and the mixtures were gently rotated for 1 hour at 4°C.

*Western Blot analyses:* Agarose beads were washed once with protein lysis buffer and three times with washing solution 1. The proteins were eluted from the beads in 27 µl of 3x SDS loading buffer [1.875 ml 1 M TRIS-HCl pH 6.8, 3 ml 20 % SDS, 3 ml glycerol, 3 mg bromophenol blue, 2.125 ml dH_2_O] with 3 µl of 1.25 M dithiothreitol (DTT). The lysate was either stored at −80°C or used directly for subsequent analyses by SDS PAGE (10%) and Western blot.

*For IP-MS*, the agarose beads were washed once with protein lysis buffer, three times with wash solution 1 and twice with washing solution 2 [150 mM NaCl, 50 mM TRIS-HCl pH 7.5, in ddH_2_O], with the beads being transferred to a fresh Eppendorf tube in the last step. The proteins were eluted from the beads in 60 µl of 6 M urea and 50 mM Tris-HCl pH 8.5. The lysate was then processed directly to mass spectrometry.

### Mass spectrometry

To obtain a comprehensive overview of the Pzg proteome at the protein level, GFP-Trap^®^ precipitates from lysed Pzg-GFP ovaries were analyzed by mass spectrometry. Precipitates (BCs) obtained with a binding control were used as negative controls. A total of three replicates, each with one IP and one BC approach, were analyzed. The analysis was performed by UHPLC-ESI-MS/MS at the Core Facility Hohenheim as follows:

First, the peptides digested overnight with trypsin were separated according to size using ultra-high-performance liquid chromatography (UHPLC) in the UltiMate 3000 RSLCnano system (Thermo Fisher Scientific) and ionised using electrospray (ESI). The signals were then recorded on the Orbitrap Exploris^™^ 480 mass spectrometer (Thermo Fisher Scientific) with subsequent Fourier transformation to determine the mass-to-charge ratio. Tandem mass spectrometry (MS/MS) was performed, in which MS1 spectra with total peptide masses were generated alongside MS2 spectra with the masses of fragmented peptides.

The spectra were initially evaluated in MaxQuant (2.6.7.0) and the integrated search engine Andromeda according to Chavan *et al*. (2025). The detected peptides were assigned to individual proteins or protein groups, which were then quantified based on the calculation of LFQ intensities. Unless otherwise specified, the MaxQuant parameters were left in default mode. The reference proteome of *Drosophila* (ID: UP000000803) was obtained on 8 October 2024 from https://www.uniprot.org (The UniProt Consortium *et al*., 2025). Only peptides with a maximum of two undigested trypsin cleavage sites and exclusively unmodified amino acids, with the exception of oxidation, carbamidomethylation and deamidation (N-terminal) modifications, were included. MS and MS/MS mass tolerance was set to 20 ppm. Peptides that could be assigned to either a single protein or multiple proteins (unique and razor) were quantified. In the latter case, a peptide was assigned to the protein that had the highest probability of actually being present in the sample. Only proteins with a false discovery rate (FDR) of 1% or less were evaluated.

The proteins identified by MaxQuant were then filtered together with their calculated LFQ intensities in Perseus (2.0.11) and statistically evaluated. Proteins that were identified solely on the basis of one or more peptides with modified amino acids, potential contaminations and reverse sequences were filtered out. In addition, a log_2_ transformation of the LFQ intensities was performed. The three IP and BC replicates were grouped and only proteins identified in all IP replicates were retained. If intensity values for proteins identified in all IP samples were missing in the BC replicates, these were supplemented by normal distribution. Significant differences between the average LFQ intensities of the IP and BC approaches were determined using a Student’s t-test.

The differences (fold change, FC) between IP and BC calculated by Perseus and the adjusted p-values in negative decimal logarithm were used to create the volcano plots. The volcano plots were created in R (4.5.1) using the packages dplyr (1.1.4) and ggplot2 (3.5.2).

Classification of proteins into cohorts (i.e. general, signalling, Chromatin, PTGS, gametogenesis, stress and apoptosis, immune, others) was based on GO for biological processes, cellular component or biological function as listed in Table S1. The mass spectrometry proteomics data have been deposited in the ProteomeXchange Consortium via the PRIDE partner repository (Perez-Riverol *et al*., 2025) with the dataset identifier PXD075330 and 10.6019/PXD075330.

### Chromatin-immunoprecipitation (X-ChIP)

ChIP experiments followed by sequencing (ChIP-seq) were used to identify the binding sites of Pzg in the genome of ovaries.

*Preparation of chromatin:* For each IP, 100 ovaries from 3 to 5 days old Pzg-GFP females were prepared in PBS. Crosslinking was then performed in 500 µl of 1% formaldehyde for exactly 10 min at RT, which was stopped by adding 500 µl 50 mM glycine. After five minutes of gentle rotation at RT, the samples were placed on ice. The ovaries were then washed three times with PBS and twice with Farnham buffer (5 mM HEPES pH 8.0, 85 mM KCl, 0.5 % NP-40, 1 tablet of protease inhibitor cocktail, 10 mM NaF, 0.2 mM Na_3_VO_4_ in ddH_2_O) for 5 minutes each at 4°C. The ovaries were then homogenized in 100 µl RIPA-2 buffer [20 mM Tris-HCl pH 7.5, 150 mM NaCl, 1 % sodium deoxycholate (SAC), 0.1 % SDS, 1 tablet of protease inhibitor cocktail, 10 mM NaF, 0.2 mM Na_3_VO_4_ in ddH_2_O] in a glass homogenizer with a size B pestle on ice and sonicated in 12 cycles (30s on, 90s off) with a Branson Sonifier Cell Disruptor B15. After centrifugation at 20.000 x g for 10 min at 4°C, the lysate was transferred to a fresh Eppendorf tube and either used directly for immunoprecipitation or stored at −80°C.

*Immunoprecipitation and purification of DNA:* For the IPs, 100 µl of lysate from ovaries were used. Before adding the antibody, 5% input control was removed from each lysate and stored at 4°C until further processing. Incubation with 10 µg rabbit ⍺-GFP antibody was carried out overnight at 4°C. The following day, 50 µl of Dynabeads^™^ Protein G were washed three times with PBS per batch. The lysate-antibody mixture was then added to the Dynabeads^™^ and incubated for 5 hours at 4°C. While the Dynabeads^™^ were washed five times for 10 minutes at 4°C, the input control was digested for 1 hour at 37°C with 1 µl 10 mg/ml RNase. After adding 713 µl PK buffer [200 mM TRIS-HCl pH 7.4, 25 mM EDTA, 300 mM NaCl, 2 % SDS, 100 µg Proteinase K] and 277 µl ddH_2_O to the input and 713 µl PK buffer and 267 µl ddH_2_O to the Dynabeads™, the samples were incubated for 3 hours in a 55°C water bath. Reverse crosslinking was performed overnight in an oven at 65°C. The following day, the DNA was extracted using phenol-chloroform, precipitated with ethanol and dissolved in 10 µl TE. The DNA concentration of the ChIP samples was determined using the Qubit 4 fluorometer (Thermo Fisher Scientific) and the Qubit 1X dsDNA HS Assay Kit. DNA fragmentation was analyzed using the 4150 TapeStation system (Agilent technologies) together with the TapeStation Analysis Software (4.1.1). The samples were stored at −20°C.

*Analysis using DNA sequencing (ChIP-seq):* The sequencing of the Pzg-ChIPs was performed by the Genomics Core Facility at EMBL Heidelberg. All samples were sequenced using Illumina’s NextSeq 500 System and NextSeq 500/550 Mid-Output v2.5 Kit, generating single-end reads with a length of 150 bp.

*Quality analysis and filtering of raw data:* To get an initial impression of the quality of the ChIP-seq samples, fingerprint plots were created using deepTools (3.5.6; tool: plotFingerprint). FastQC (0.12.1) was also used to determine the number and quality of the reads. The ChIP-seq raw data were first subjected to quality control and trimming in Trim Galore (0.6.10; setting: --quality 20), whereby adapter sequences and all bases with a Phred quality score below 20 were removed. The sequences were then filtered for low-complexity regions using BBMap (39.19; tool: BBduk.sh, settings: entropy=0.35 entropywindow=18 entropyk=4), using an entropy threshold of 0.35. To remove mitochondrial sequences, the reads were aligned to the corresponding sequences (FlyBase dm6 genome release 6.59, ChrM) using Bowtie2 (2.5.4) and only non-aligned reads were used. To determine the quality and number of filtered reads, a further analysis was performed using FastQC (0.12.1).

*Alignment to the Drosophila genome:* The sequence for the *Drosophila* genome dm6 (release 6.59) was obtained from FlyBase (https://flybase.org, release FB2024_04). Bowtie2 (2.5.4; settings: -k 1, --very-sensitive) was used for alignment. The generated alignments in Sam format were then converted to Bam files and sorted using Samtools (1.21; tools: view, sort). Duplicates were removed with Picard (3.3.0; tool: MarkDuplicates) and blacklist regions (Amemiya *et al*., 2019) were removed by intersection with Bedtools (2.31.0; tool: intersect).

*Normalisation and visualisation of alignments*: The final alignments were converted into Bigwig files for visualisation and quantification using deepTools (3.5.6; tool: bamCoverage; setting: --normaliseUsing RPKM) and normalised to the number of reads. The enrichments of the IPs were then calculated in comparison to the respective input using deepTools (3.5.6; tool: bigwigCompare; settings: --pseudocount 1 --operation log2) and averaged across the three replicates using deepTools (3.5.6; tool: bigwigAverage). The alignments were visualised in IGV (2.18.4).

Data have been deposited to Gene Expression Omnibus (GEO) database of the National Library of Medicine (NCBI, Bethesda, US) under the accession number GSE324254.

### Quantitative RT-PCR

For each reaction four biological and two technical replicates were performed. All used materials are listed in the Key Resources Table.

*Transposon transcripts:* Relative mRNA levels of several transposon transcripts and the Notch antagonist *Tom* was determined by performing quantitative RT-PCR (qRT-PCR) on four biological and two technical replicates of 25 ovaries each from freshly hatched females of the genotype *nos*-Gal4::UAS-shRNA-*w* (reference/control) and *nos*-Gal4::UAS-shRNA-*pzg.* Poly(A)^+^ RNA extraction, cDNA synthesis and real time qPCR was performed as described before (Kober *et al*., 2019) using the MIC magnetic induction cycler. Sequences of the primers are listed in Star materials table. Relative quantification was performed with workbench version v2.0.7 based on REST, taking target efficiency into account.

*piRNA cluster transcripts:* Total RNA from ovaries of 50 freshly hatched females of the genotypes *nos*-Gal4::UAS-shRNA-*w* and *nos*-Gal4::UAS-shRNA-*pzg,* respectively, were prepared using miRNeasy Micro Kit and digested with DNase at 25°C for 15 min. cDNA synthesis was performed using ProtoScript II First strand cDNA synthesis kit, with 500 ng total RNA, using specific primers. This means that the mixture for a cDNA synthesis reaction contains specific primers for all reference genes and for either the plus or minus strand of 5 different clusters. The synthesis for the plus and minus strands of the clusters is carried out in separate approaches, so the same primer pair can be used for quantification for both strands in the subsequent qPCR. For each reverse transcription 6 µl total RNA (500 ng) + 2 µl 2.5 µM primer stock solution (0.25 µM final concentration/20 µl) was denaturated 5 min at 65°C, then 10 µl 2x ProtoScript II Reaction Mix and 2 µl 2x ProtoScriptII Enzyme Mix was added. cDNA synthesis was performed for 1h at 48°C, followed by 5 min inactivation at 80°C. cDNA was diluted with 50 µl nuclease-free water. RT-qPCR reactions were done as described in Kober *et al*., 2019.

The generally expressed genes *CPSF*, *cyp33*, *SdhA* and *tbp* served as global reference, whereas *dlp* and *Lamin C* were selected as specific reference genes for somatic TEs or the *flam* piRNA cluster, based on their expression predominantly in the somatic cells of the germarium.

## Results

### Nurf- and Dref-independent function of *pzg* in the germline

Previously, we demonstrated that *pzg* is intimately involved in germ cell survival and the well-balanced stem cell–niche communication in the female germline. When *pzg* activity was specifically downregulated in the germline, GSC’s failed to differentiate and instead accumulated in the germarium outside the niche to eventually being eliminated by cell death. Consequently, only atrophied, rudimentary ovaries developed, and the females were completely sterile (Figure 1A,A’, Kober *et al.,* 2019). The underlying cause, however, remained unresolved so far. In larval somatic tissues, Pzg has been linked to epigenetic regulation, for example by its association with the nucleosome remodelling NURF complex, thereby influencing various signalling cascades (Kugler and Nagel, 2010; Kugler *et al*., 2011). A germline specific knockdown of *Nurf301*, a central and exclusive NURF subunit, resulted in young adult females with morphologically inconspicuous ovaries. These harboured the wildtype complement of 2-3 GSCs in the stem cell niche at the apical tip of the germarium (Figure 1B,B’). This finding agrees with earlier reports, whereby *Nurf301* mutants display premature/precocious GSC differentiation in the female and male germline, thus ensuring their maintenance in the niche, which is in contrast to *pzg* knock down phenotypes (Ables and Drummond-Barbosa, 2010; Cherry and Matunis, 2010). In addition, Pzg protein was also identified as an integral part of the Trf2/Dref complex, involved in the regulation of proliferation related genes (Hochheimer *et al.,* 2002; Kugler and Nagel, 2007). Thus, loss of *pzg* activity in larval tissues results in a disruption of cell cycle progression and consequently in growth defects. Although maternal depletion of *Dref*, which targets the complex to DRE-binding sites on DNA, caused atrophied ovaries that superficially resembled those observed after *pzg* depletion (compare Figure 1A with C), no accumulation of undifferentiated GSCs was observed (Figure 1C’). Moreover, no specific signs for a cell cycle arrest in either the S phase (revealed by EdU incorporation) - or in M phase (indicated by PH3 staining) was observed in *pzg-*deficient GSCs, as we would have expected if loss of *Dref* activity were causal (Figure 1D,E). These findings indicate that the effects of *pzg* loss in the female germline are not due to compromised Nurf- and/or Dref function, raising the question how Pzg might exert its influence in this developmental context.

**Figure 1.**
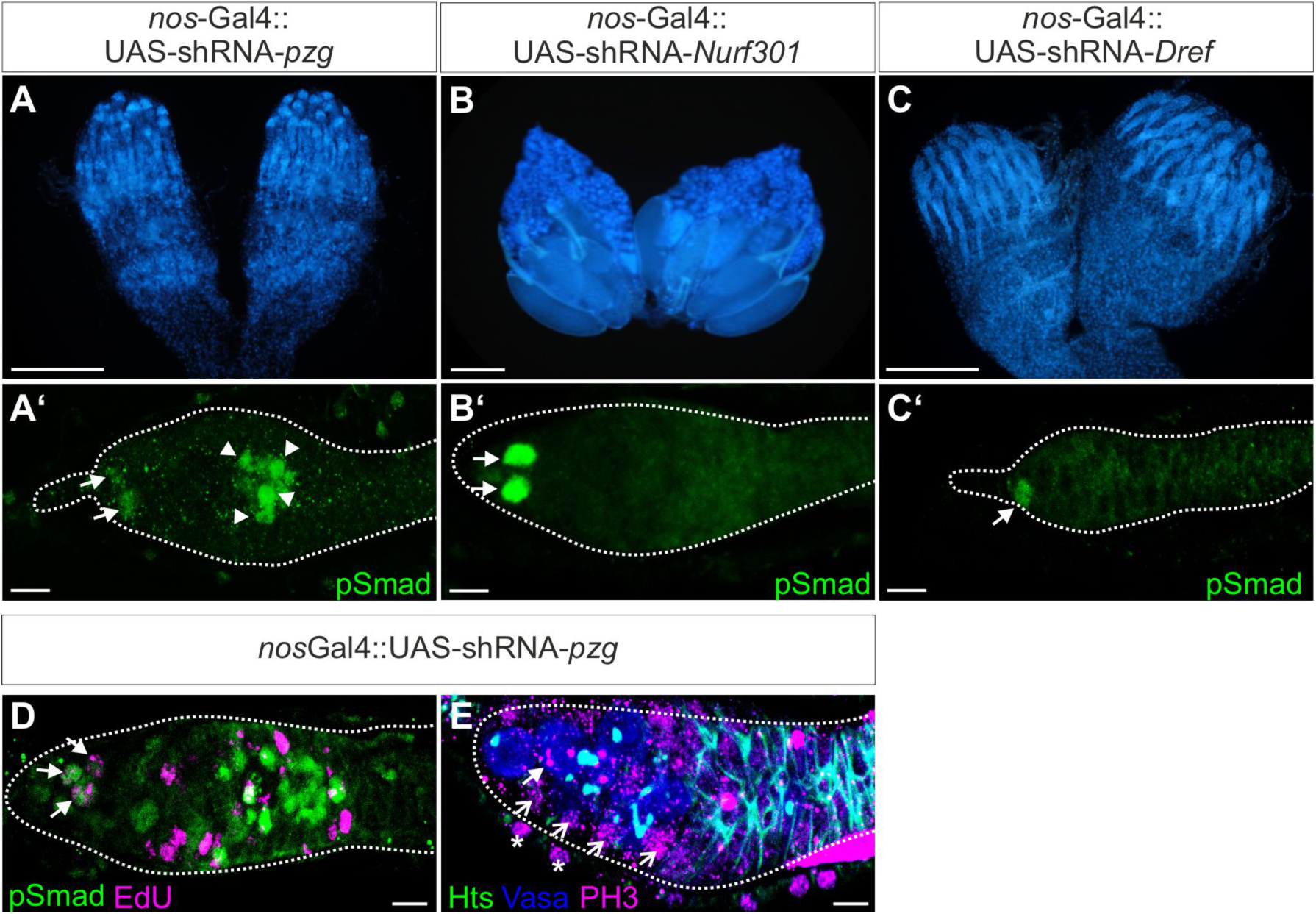
Loss of pzg in the germline does not compare to loss of Dref or Nurf301. (**A-C**) Ovaries stained with DAPI after depletion of *pzg* (A), *Nurf301* (B) or *Dref* (C) via UAS-shRNAi induction during germ cell development using *nos*-Gal4. Scale bars, 200 µm. (**A’-C’**) *pzg* depletion increases the number of GSCs in 0-2 days old germaria (A’), whereas a wild type complement of 2-3 GSCs is detected in *nos*-Gal4::UAS-shRNA-*Nurf301* mutant germaria (B’). Loss of *Dref* in the germline is correlated with a loss of GSCs (C’). GSCs are highlighted by a strong pSmad signal (green, arrows). Scale bars, 10 µm. (**D,E**) Dividing GSCs and germline cells are detected in *nos*-Gal4::UAS-shRNA-*pzg* mutant germaria. Scale bars, 10 µm. (D) Co-staining of pSmad (green) and EdU (magenta); note overlap in GSCs (arrows). (E) Triple-staining with ⍺-Vasa (blue) and ⍺-Hts (green, seen as turquoise in the overlap) with ⍺-PH3 (magenta); note labelled GSC (closed arrow) and germline cells (open arrows). Asterisks mark cells of the outer sheath of the ovariole.

### Increased genomic instability in *pzg* mutant cells

Reports on signs of moderate telomere instability in the *pzg* mutants (Silva-Sousa *et al.,* 2012) prompted us to assess DNA integrity in *pzg^66^* null mutants as an indicator of genomic instability in *pzg* depleted cells. Larval development of *pzg^66^* homozygotes is significantly delayed and ends in the second instar (Kugler *et al*., 2011). Thus, we measured the incidence of DNA double strand breaks in freshly hatched first instar larvae, before apoptotic processes occur, as not to confuse cause and effect of *pzg* depletion. One of the first responses to DNA double strand breaks is the phosphorylation of the histone variant H2Av (g-H2Av in *Drosophila*), which serves as a platform for proteins of the repair machinery and can be detected there until repair of the DNA damage is completed (summarized in Bekker-Jensen and Mailand, 2010; Polo and Jackson, 2011). Immunoblots from nuclear extracts uncovered a robust signal of this histone modification in the *pzg^66^*deficient larvae, which was not present at this stage in the wildtype control (Figure 2A). In the *Drosophila* female germline, DNA double strand breaks are specifically induced during meiotic recombination, which can be detected in a wildtype germarium in region 2a/b, but never in GSCs to avoid the transmission of incorrect/defective genetic information to the next generation (Figure 2B-B’). However, if we specifically downregulated *pzg* activity in the germline, a strong g-H2Av signal was detected in *pzg* depleted GSCs, identified by the co-staining of the GSC specific marker phospho-Smad (pSmad) (Figure 2C-C’). DNA damage was observed in GSCs located in the anterior stem cell niche as well as in those accumulated further posteriorly. Consistent with the incidence of DNA damage in *pzg* deprived GSCs, p53 biosensor activity (p53R-GFP) was also triggered, known to be selectively sparked as a safeguard in GSCs when genome destabilizing stress occurs in the germline (Figure 2D) (Wylie *et al*., 2014; Wylie *et al*., 2016). Thus, *pzg* activity is indispensable to protect genome integrity in the cells of the female germline.

**Figure 2.**
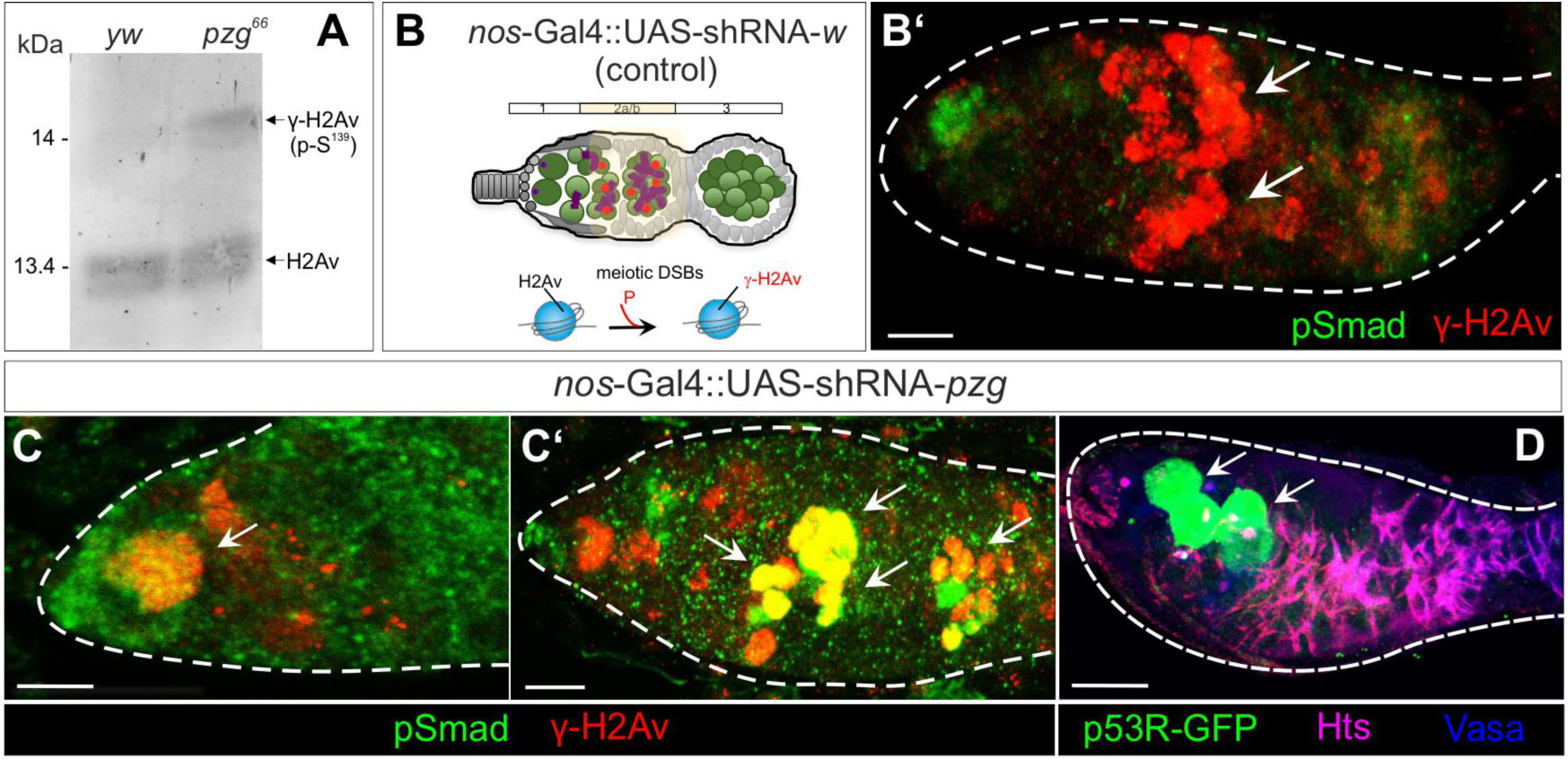
Loss of pzg in the germline causes genomic instability. (**A**) Immunoblot with nuclear extracts from *pzg*^6*6*^ mutant versus *y^1^ w^67c23^*control (*yw*) early larvae, detecting γ-H2Av phosphorylated at Serine^139^ (by using an γ-H2AX antibody) versus H2Av. (**B**) Schematic drawing of a germarium with the developmental stages 1, 2a/b and 3. GSCs (green) are characterized by a spherical spectrosome (purple dot), whereas developing cysts and cystoblasts (green) are characterized by branched fusomes (purple bar). Meiotic recombination occurs in the region 2a/b, highlighted in light yellow, resulting in double-strand breaks in the DNA (DSBs) and phosphorylation (P) of the histone variant H2Av (red dots). (**B’)** In the control (*nos*-Gal4::UAS-shRNA-*w*) γ-H2Av signals (red) accumulate in region 2a/b (arrows). GSCs (green, pSmad) remain unlabelled. (**C-C’**) After downregulation of *pzg* (*nos*-Gal4::UAS-shRNA-*pzg*) γ-H2Av foci (red) are detected in pSmad positive GSCs (green) in the niche (C), as well as outside the niche (C’) (arrows, overlay appears yellow). (**D**) Germline depletion of *pzg* causes p53R-GFP reporter activity in GSCs (green, arrows) (blue: Vasa, magenta: Hts). Scale bars in all panels, 10 µm.

### The DNA damage network is disturbed in *pzg* depleted germ cells

DNA damage induces stress response pathways that are also used in other instances of environmental stress, notably the JNK pathway, and may be mediated by miRNAs acting as systemic signaling components (Figure 3A) (reviewed in Luhur *et al*., 2013). In human and zebrafish, for example, miR-125b ensures low p53 expression levels. This repression is relieved in response to DNA damage resulting in p53-mediated apoptosis (Le *et al.,* 2009; Zhang *et al.,* 2009; reviewed in Leung and Sharp, 2010). Interestingly, previous work documented a central role for *mir-125* in the *Drosophila* stem cell niche by targeting *Twin of m4* (*Tom*), a negative regulator of the Notch signalling pathway (Bardin and Schweisguth, 2006; Yatsenko and Shcherbata, 2018). Accordingly, we would expect the downregulation of Notch activity in female germaria and the upregulation of *Tom* mRNA levels upon *pzg* depletion. Indeed, the expression of a Notch Response Element reporter fused to eGFP (*NRE-eGFP*) was completely lost from the stem cell niche in young adult germaria (0-3d) depleted of *pzg* (Figure 3B-C’’). Moreover, qRT-PCR analyses indicated an app. 3.5-fold increase in *Tom* mRNA levels upon *pzg* knock down (Figure 3D).

**Figure 3:**
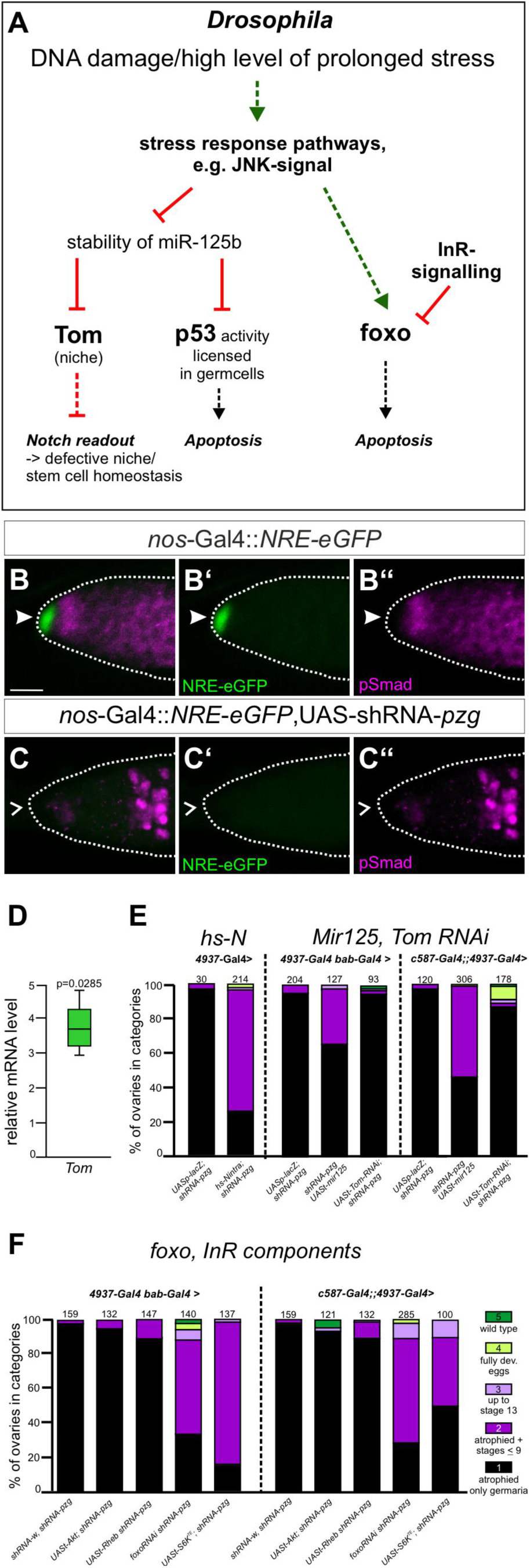
The DDR network influences the pzg mutant germline phenotype. (**A**) Schematic illustrating the network of the main players and factors involved in DNA damage response (DDR) and prolonged stress. Stress response is coordinated by the JNK-signaling pathway, mediating apoptosis via the InR/foxo axis and p53 activity downstream, as well as a balanced Notch activity at the niche/stem cell interface. (**B-B’’**) Notch signalling activity, monitored by the NRE-eGFP reporter (green) in the germline niche (arrowhead) in the control (*nos*-Gal4::NRE-eGFP); germ cells labelled magenta (⍺-pSmad). (**C-C’’**) Notch activity, i.e. NRE-eGFP expression (green) in the germline is lost upon downregulation of *pzg* (*nos*-Gal4::NRE-eGFP, UAS-shRNA-*pzg*). Scale bar, 10 µm for (B-C’’). (**D**) In comparison to the control, *Tom* transcript levels are significantly increased in *nos*-Gal4::UAS-shRNA-*pzg* mutant germaria. Four biological and two technical replicates were performed. Median corresponds to expression ratio; mini-max depicts 95% confidence. Expression ratio was significant at the level of <0.05 using PFRR from REST: p=0.0285. (**E-F**) Rescue-assays of atrophied ovaries by changing the level of Notch signalling components (E) or *foxo* and members of the InR-pathway (F) in the cells of the niche (using *bab1*-Gal4) or in escort cells (with *c587*-Gal4) in *pzg* deficient germaria, as indicated. Ovaries of 3-5 days old females were classified in 5 different categories (each differently colored in the bar chart) according to their state of development: **1 (black)**: atrophied ovaries, only germaria-like structures; **2 (dark purple)**: rudimentary ovaries with some follicles developed up to stage 9; **3 (light purple)**: development up to stage 13 in some ovarioles; **4 (light green)**: occasional complete egg development; **5 (dark green)**: comparable to a wild type ovariole. Number of analyzed ovaries is given above the bars; the examined genotype is given below. Increasing Notch (E) and InR (F) signalling, respectively, or reducing *foxo* activity (F) in the niche and/or escort cells ameliorated the atrophied germline phenotype caused by *pzg* knock down in the germline.

According to our expectations, a subtle overexpression of Notch rescued the atrophic ovary phenotype of *shRNA-pzg* mutants (Figure 3E). Since the overexpression of a constitutively activated Notch receptor is often associated with pleiotropic phenotypes, we opted for a heat-inducible Notch construct (Lieber *et al.,* 1993). Starting in early larval stages, one daily heat shock (1h 37°C) was applied until flies hatched from which the ovaries were examined for ovariole development. In contrast to the untreated *shRNA-pzg* mutants, females that experienced continuous Notch supplement developed eggs up to stage 9 in approximately 80% of the ovaries analyzed. Occasionally, even wildtype ovarioles were obtained (Figure 3E). Moreover, a specific overexpression of *mir-125* or the depletion of *Tom* in either niche cells (using *bab*-Gal4) or in escort cells (using *c587*-Gal4) also improved the development of *pzg* depleted ovaries quite well (Figure 3E).

Foxo is another prominent player in stress resistance and is supposed to preserve cellular homeostasis and resilience and mediate longevity in vertebrates as well as in *Drosophila* (reviewed in Martins *et al*., 2016; Rodriguez-Colman *et al*., 2024) (Figure 3A). Apart from its well-established role as a key transcriptional regulator of the InR-pathway, Foxo is also navigated by the stress-responsive JNK-signalling pathway (Wang *et al.,* 2005; Puig and Mattila, 2011; Martins *et al.,* 2016). In agreement with our earlier observation of JNK-activation in *pzg-*depleted germline cells (Kober *et al.,* 2019), we observed a strong rescue of the *shRNA-pzg* mutant phenotype by the depletion of *foxo* in the germline: Only 34% of the analyzed ovaries were still atrophied, whereas 55% developed until stage 9, 6% up to stage 13 and 2% had wildtype morphology (Figure 3F). An even slightly better rescue was observed with a RNAi-mediated *foxo* reduction in escort cells, whereby 72% of the ovaries developed at least until stage 9 (Figure 3F). However, when we overexpressed the upstream kinase Akt, known to inhibit Foxo activity, or the InR-signalling pathway members Rheb or S6K, the rescue effects were far less pronounced, indicating that genomic stress rather than failure of InR signalling activity or growth defects are the major consequences of a *pzg* depletion in the germline.

### Deregulated transposon activity in *pzg* depleted germline and somatic tissue

Transposable elements (TEs) pose one of the biggest threats for genome integrity, and cause *p53* activity selectively in germline stem cells, indicating that p53 functions to restrain TE activity (Wylie *et al.,* 2014; Wylie *et al.,* 2016). Thus, the strong p53 biosensor activity in *pzg* deficient germline cells may result from a de-repression of TEs. To test this idea, we first made use of two established TE-reporters (sensors) in the germline, *burdock-lacZ* and *Het-A-lacZ* as well as *gypsy-lacZ*, as an example of a somatic transposon reporter (Handler *et al*., 2013; Czech *et al*., 2013). RNAi-mediated depletion of *pzg* in cells of the germline resulted in strong *ß-Gal* signals, i.e. *Het-A* and *burdock* sensor de-repression, however, no increased expression of *gypsy-lacZ* in comparison to the control (Figure 4A), indicating that loss of *pzg* activity favours the de-repression of germline specific TEs. To extend these findings, we examined the expression levels of several TE transcripts and detected an increase in the range of 1.6-fold (*Hobo*) up to 41-fold (*R2*) (Figure 4B). In general, TEs expressed in the germline belonging to Non-LTR, LTR and DNA family of transposons were de-repressed, whereas *gypsy* and *zam*, both members of the somatic LTR family, remained unchanged (Figure 4B’). As the strongest de-regulation was observed for retrotransposons [e.g. R2 (non-LTR) and 297 (LTR)] that employ an RNA intermediate during mobilization, we asked whether suppression of reverse transcription might mitigate the atrophied ovariole phenotype of *pzg* knock down mutants, by allowing the animals to develop on diet with yeast paste enriched with lamivudine (3TC), a clinically used nucleoside reverse transcriptase inhibitor (Lin *et al*., 2020). Although presumably quite inefficient due to oral application, still 40% of the analyzed ovaries developed until stage 9, demonstrating that de-repression of retrotransposons contributes to the *shRNA-pzg* mutant germline phenotype (Figure S1).

**Figure 4:**
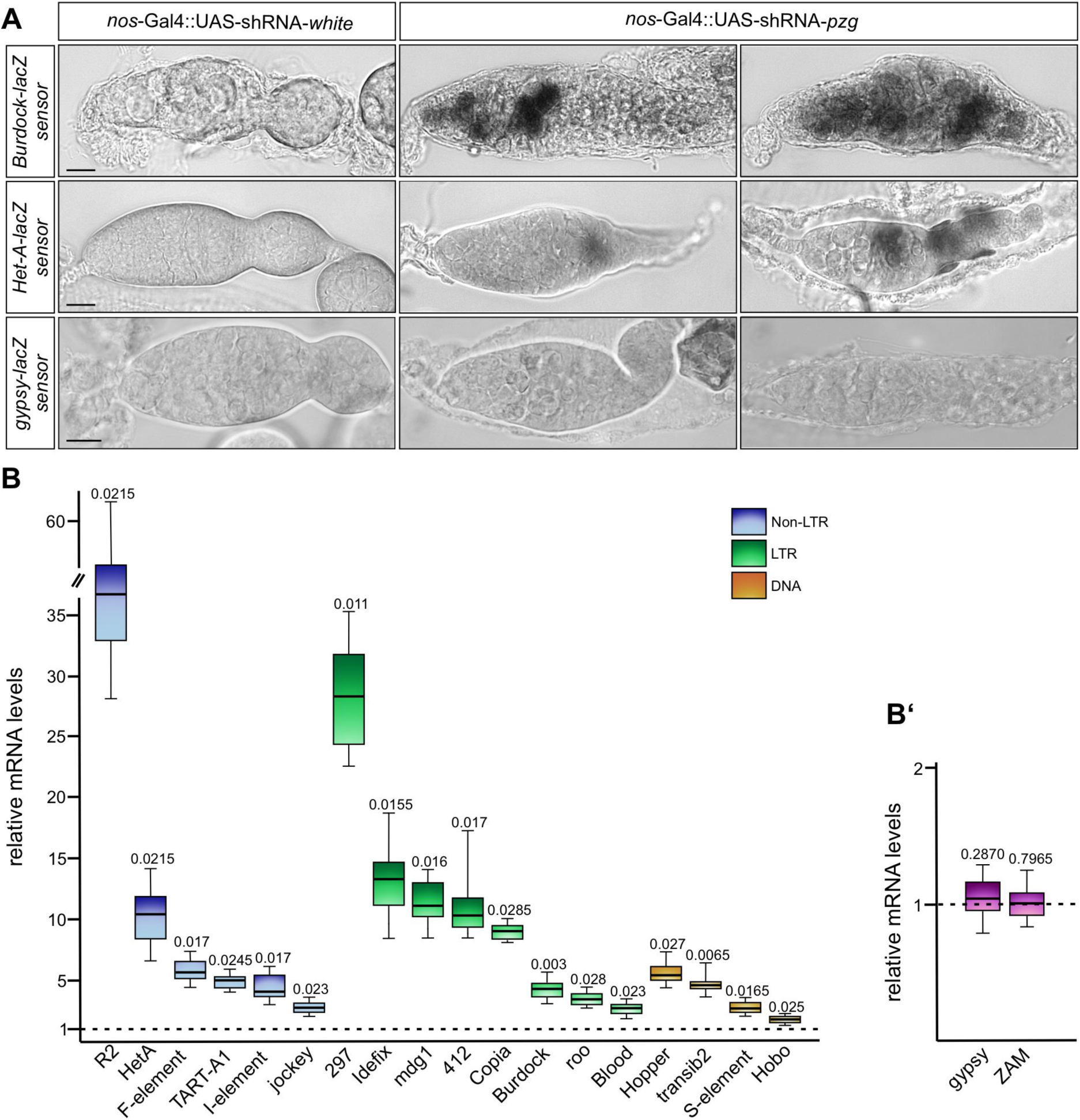
De-repression of transposons in pzg mutant background. (**A**) Depicted are beta-galactosidase-stainings of germaria expressing either *burdock*-lacZ sensor (upper row), *Het-A*-lacZ sensor (middle row) or *gypsy*-lacZ sensor (lower row) in control ovaries (*nos*-Gal4::UAS-shRNA-*w*, left) and *pzg* mutant ovaries (*nos*-Gal4::UAS-shRNA-*pzg*, middle and right). Whereas no transposon activity is detected in the control, both *burdock* and *Het-A* transposons, but not the somatic *gypsy* transposon, are de-repressed in *pzg* mutant germaria. Scale bars, 10 µm. (**B,B’**) Quantitative RT-PCR of germline (B) and soma (B’) transposon RNA levels from ovaries of freshly hatched *nos*-Gal4::UAS-shRNA-*pzg* females relative to the control (*nos*-Gal4::UAS-shRNA-*w*). (B) The steady state RNA level of various transposons expressed in the germline and belonging to different families [Non LTR (blue), LTR (green) and DNA (brown)] is significantly increased to various degrees. *CPSF*, *cyp33*, *SdhA* and *tbp* were taken as references for determining relative quantities by REST. (B’) In contrast, *gypsy* and *ZAM* transcripts derived from two somatic transposons, are unchanged. Here, *LamC* and *dlp*, expressed in the soma, were taken as reference. (B,B’) Four biological and two technical replicates were performed. Median corresponds to the expression ratio; mini-max depicts 95% confidence. All expression ratios shown in (B) were significant at the level of p<0.05 using PFRR from REST. Exact p values are given above the bars.

### Pzg is involved in piRNA precursor expression

In the germline, TE transposition is kept in check with the help of the Piwi-interacting RNAs, the piRNAs (reviewed in Czech *et al*., 2018; Ozata *et al*., 2019). piRNAs are encoded in genomic piRNA clusters (Brennecke *et al*., 2007; Senti and Brennecke, 2010). The uni-directional cluster *flamenco* (*flam*) is active only in the soma, in contrast to the uni-strand cluster *20A1*, which is active in the soma and germline. The three major dual-strand clusters, located at cytological position *38C1*/*38C2*, *42AB* and *80F*, are transcribed in germline cells only (Brennecke *et al.,* 2007; Senti and Brennecke, 2010; Mohn *et al.,* 2014). Transcription of the latter occurs in a non-canonical way, neither using classical transcriptional start sites, i.e. promoters, nor transcription termination (reviewed in Czech *et al.,* 2018; Ozata *et al*., 2019). Instead, transcription of the dual-strand piRNA clusters is mediated by the Rhino/Deadlock/Cutoff complex (RDC complex), which further recruits the Moonshiner/Trf2/TFIIA-S complex to initiate transcription from both strands in a promoter independent manner (Andersen *et al.,* 2017). As Pzg was originally identified as integral part of the Trf2/DREF transcription-initiation protein complex (Hochheimer *et al*., 2002), we wondered whether Pzg might be involved in the initiation of piRNA-precursor transcription. In this case, we would expect lowered piRNA levels from these bi-directional clusters in the *pzg*-depleted germline. To this end, we performed qRT-PCR analyses to quantify the piRNA precursor levels from both strands in *nos*-Gal4::UAS-shRNA*-pzg* compared to *nos*-Gal4::UAS-shRNA*-w* control ovaries. Indeed, expression of the somatic *flamenco* locus was indistinguishable from the control, whereas that of *20A1* was reduced by half (Figure 5A), in agreement with its activity in both soma and germline cells (Brennecke *et al*., 2007; Senti and Brennecke, 2010). In contrast, piRNA precursor transcription was considerably reduced (>95%) from either strand in the RDC-dependent clusters *42AB* and *80F*, respectively (Figure 5A), demonstrating the requirement of Pzg for RDC-mediated piRNA precursor generation. Intriguingly, transcription of both the *38C1* and *38C2* was differentially affected. *38C1* expression was down to about 25-50% with a notable difference between the plus and the minus strand. The latter compared to the expression of the uni-transcribed cluster *20A1* (Figure 5A). And whilst *38C2* plus strand expression was reduced to nearly 10% of the control, the minus strand showed a roughly 3-fold increase. Increased piRNA transcription from the *38C1* cluster was observed in *moonshiner* mutants by use of adjacent promoters. Moreover, a minus-strand bias was also observed upon a plus-strand promoter deletion in the *38C1* cluster (Andersen *et al.,* 2017). We conclude that Pzg supports the promoter-driven cluster transcription as well.

**Figure 5:**
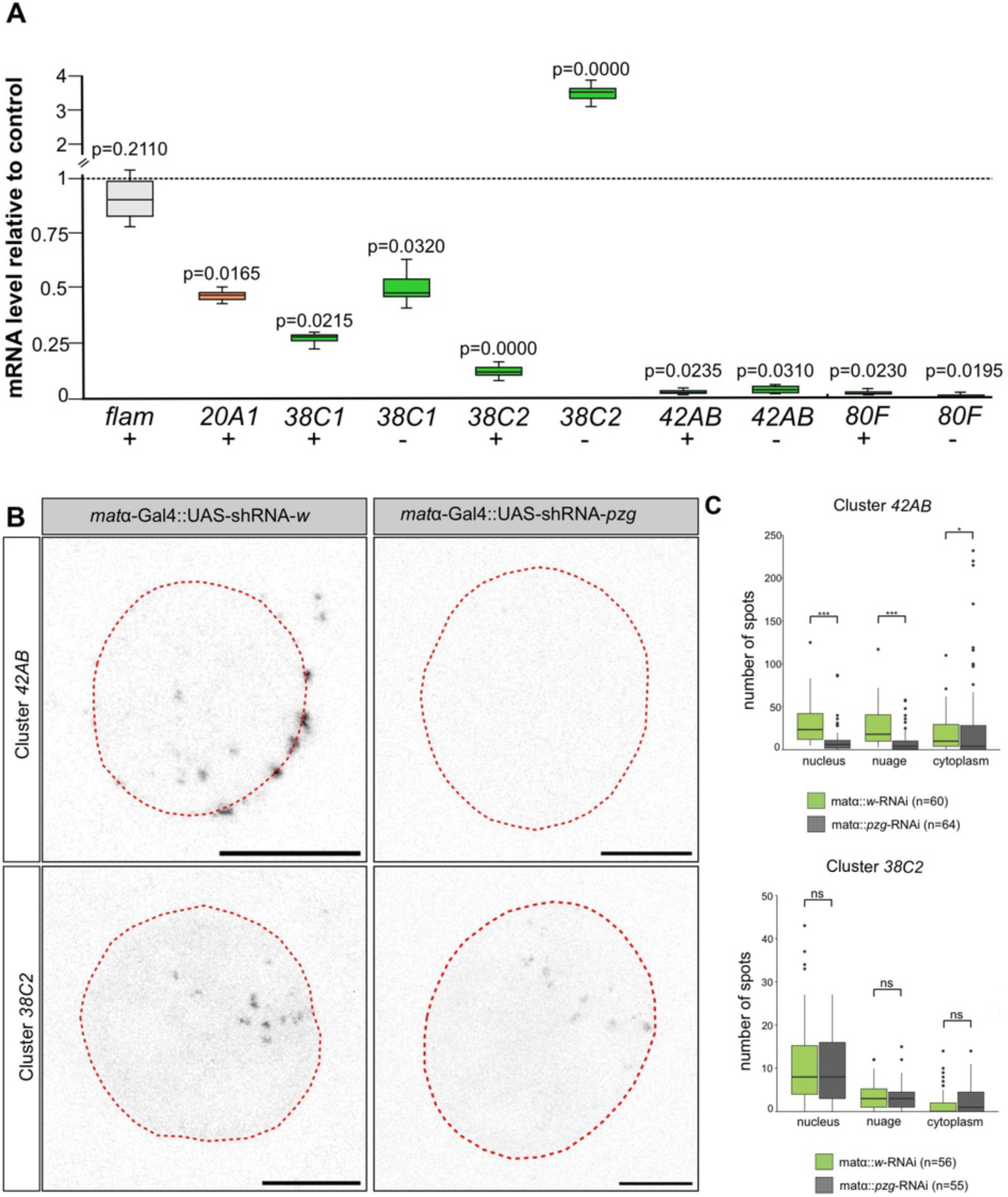
piRNA dual-strand cluster transcription is changed in pzg mutant ovaries. (**A**) Strand-specific qRT-PCR for RNA derived from the uni-strand somatic piRNA cluster *flam*, soma- and germline expressed uni-strand cluster *20A1* and the germline-expressed dual-strand clusters *38C1/C2*, *42AB* and *80F*, compared between ovaries from the control (*nos*-Gal4::UAS-shRNA-*w*) and upon *pzg* knock down in germline cells (*nos*-Gal4::UAS-shRNA-*pzg)*. Whereas RNA levels from the somatic cluster *flam* were unchanged, RNAs from cluster *20A1* and from both strands of clusters *38C1*, *42AB* and *80F* were starkly reduced in *pzg* depleted germline cells. RNA from the plus strand of *38C2* was reduced as well, whereas RNA levels derived from the *38C2* minus strand were increased more than threefold. Four biological and two technical replicates were performed. Median corresponds to the expression ratio; mini-max depicts 95% confidence. All expression ratios shown are significant at the level of p<0.05 using PFRR from REST. Exact p values are given above the bars. The ubiquitously expressed genes *CPSF*, *cyp33*, *SdhA* and *tbp* were taken as references for determining relative quantities by REST of germline expressed clusters *20A*, *38C1/C2*, *42AB* and *80F*, whereas *LamC* and *dlp*, both expressed somatically, were taken as reference for somatic cluster *flam*. (**B**) RNA-FISH signal detection of precursor transcripts derived from piRNA cluster *42AB* and *38C2* in control (*mat⍺-*Gal4::UAS-shRNA-*w*) and *pzg*-depleted nurse cells (*mat⍺-*Gal4::UAS-shRNA-*pzg*). Dotted line: Nuclear outline based on DAPI staining. Scale bars, 10 µm. **(C)** Statistical evaluation of RNA-FISH spots in the nucleus, the nuage and the cytoplasm. n=55-64 nurse cells as indicated from at least 3 independent experiments were evaluated. Pzg deficient nurse cells have significantly less *42AB* spots in all compartments, whereas the number of *38C2* spots did not vary significantly. Mann-Whitney-Wilcoxon test. *** p<0.001; * p<0.05. Not significant (ns, p≥0.05).

We further performed RNA fluorescent *in situ* hybridization (RNA-FISH) analysis for cluster *42AB* and *38C2* transcripts in *pzg-shRNA* depleted cells of the germline. As the localization of piRNAs is best analyzed in nurse cells, we used a different Gal4 driver line, *mata-*Gal4VP16, allowing the development of later egg stages after *pzg-shRNA* induction (Figure 5B). Robust reduction of cluster *42AB* FISH signals was detected in *pzg*-depleted nurse cells with respect to foci in the nucleus (−75%), the nuage (−78%) as well as the cytoplasm (−60%), supporting reduced *42AB* piRNA expression, affecting piRNAs precursors levels also outside the nucleus (Figure 5C). In contrast, *38C2* FISH probes did not uncover significant differences in expression and localization in the analyzed nurse cells (Figure 5B,C). As the FISH probes detect both strands, reduction of plus and increase of minus strand signals might be levelled out, in line with the idea, that *38C2* transcription is compensated for on one strand.

### Pzg physically associates with the Rhino-Deadlock-Cutoff-Moonshiner complex

As *pzg*-depletion in germline cells resulted in reduced levels of piRNA precursors, we wondered whether Pzg might be associated with the alternative RDC-Moon/Trf2 transcription initiation complex. We took advantage of transgenic fly strains expressing tagged versions of either Rhino, Deadlock, Moonshiner or Trf2. We trapped these components from ovary lysates by performing immunoprecipitations and checked for co-purification of Pzg protein from these lysates. Throughout our experiments, we were able to co-purify Pzg with Moonshiner and Trf2 as well as with Rhino and Del (Figure 6A).

**Figure 6:**
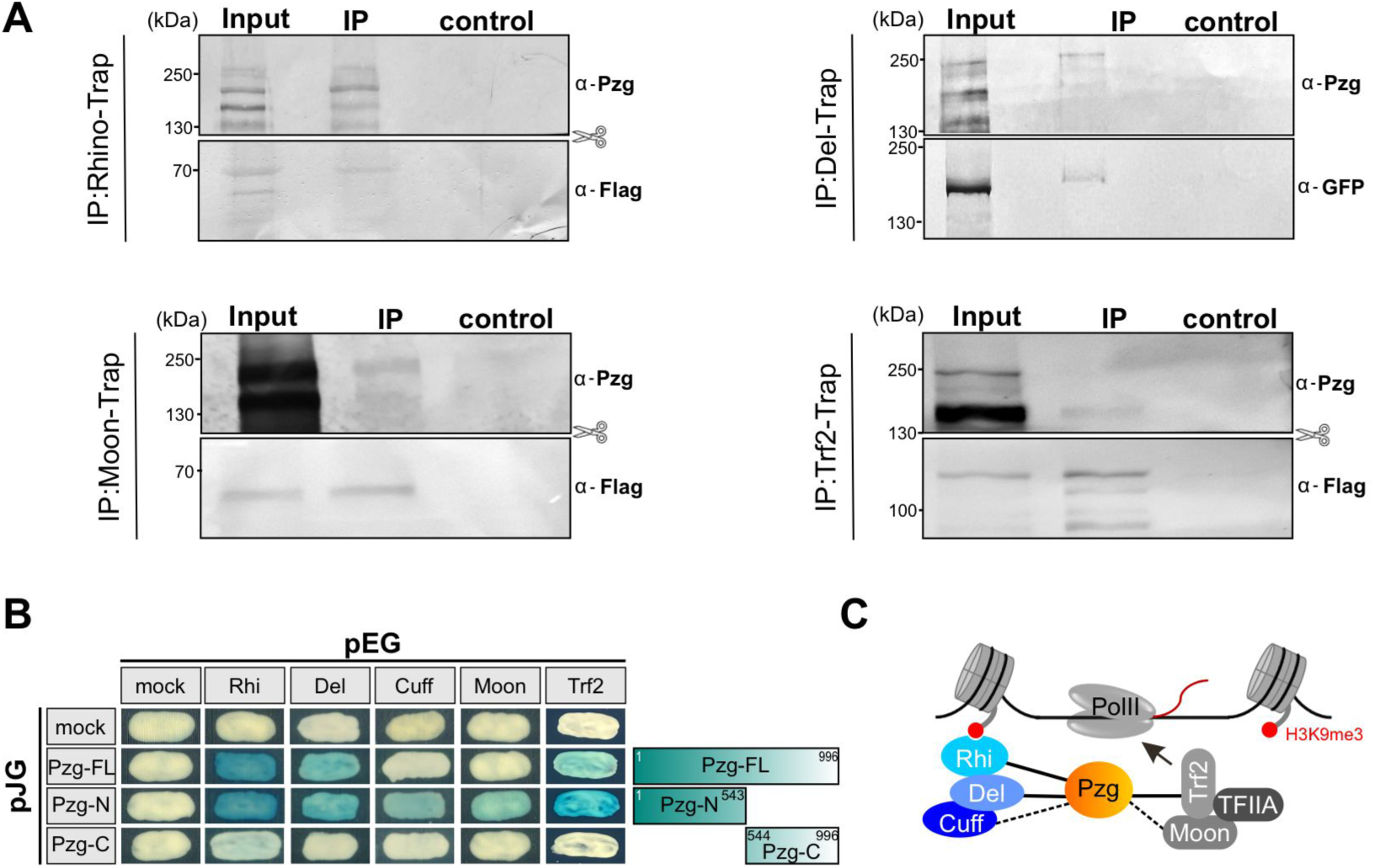
Pzg associates with members of the RDC-initiator complex. (**A**) Western Blot analysis of co-immunoprecipitation experiments: Rhino-, Del-, Moon- or Trf2-tagged proteins were trapped using magnetic GFP-Trap agarose beads. Input, 10% of IP-trap; binding control were agarose beads. For detection either GFP or Flag antibodies were used. Blot were probed with anti-Pzg antibody showing co-precipitation in all four traps. (**B**) The Brent yeast two-hybrid assay was performed to test for direct interaction between Pzg and members of the RDC complex. Rhi, Del, Cuff, Moon and Trf2 were used as bait by fusion with the DNA binding domain lexA in pEG-vector, whereas Pzg proteins (Pzg-FL, full length), N-terminus (Pzg-N, aa 1-543) and C-terminus (Pzg-C, aa 544-996) were fused to the acidic domain in the pJG vector. Positive interaction is characterized by blue yeast colonies due to the activation of the *lacZ*-reporter. Interaction was observed with the N-terminal part of Pzg, but not with the C-terminal part. (**C**) Schematic of the RDC and Moon/Trf2 complexes, including confirmed contacts to Pzg.

To date, the molecular link between RDC and the Moon/Trf2/TFIIA-S complexes is still elusive, although co-precipitation of Del and Moon indicates close contacts (Andersen *et al.,* 2017; Riedelbauch *et al.,* 2025). As Pzg is present in complexes with Trf2 (Hochheimer *et al*., 2002), Pzg may be involved in docking the transcription initiation complex to the RDC complex. In this case, we might expect direct protein-protein interactions amongst some of the components, which we probed in yeast two-hybrid assays. These studies revealed that the full-length Pzg protein robustly bound to Rhi, Del and Trf2 (Figure 6B). We could narrow down the interaction domain to the N-terminal part of Pzg, which in addition also bounds weakly to Cuff and Moon (Figure 6B). The N-terminal domain harbors the zinc-finger domains as well as a zinc-finger associated domain involved in Pzg dimerization (Eggert *et al*., 2004; Bonchuck and Georgiev, 2024). As smaller constructs tend to be expressed at higher levels, we were fortunate to uncover these weaker protein-protein interactions. The C-terminal part of Pzg did not interact with either component of the RDC-Moon/Trf2 complex (Figure 6B). We conclude that Pzg, via direct interactions with the RDC complex, promotes the recruitment of the Moon/Trf2 complex for the initiation of non-canonical transcription of dual-strand piRNA clusters, and hence is required for piRNA production in the Piwi-mediated transposon defense.

We next performed X-Chip experiments to determine the binding of Pzg at the *42AB* and *38C1/C2* loci compared to RDC complex binding. In contrast to Rhino that largely covers the entire piRNA loci, Pzg localization was more similar to Del and Cuff (Figure 7A). Pzg was enriched at the 3’ end of cluster *42AB* and at the 5’ end of the clusters *38C1* and *38C2*, overlapping with Del, and to a lesser degree with Cuff (Figure 7B). This positioning is along the lines with a role of Pzg in the initiation of transcription rather than the elongation, in agreement with our model that Pzg links the Moon/Trf2 initiation complex to RDC. Within the *42AB* complex, we noted an alternate pattern of Pzg binding, reflecting its near-complete absence from the annotated TEs with interjacent presence. A similar binding was observed for Del and Cuff, perhaps reflecting the sites of RDC-complex formation, whereas Rhi spread more equally in agreement with its role in heterochromatin formation (Figure 7C).

**Figure 7:**
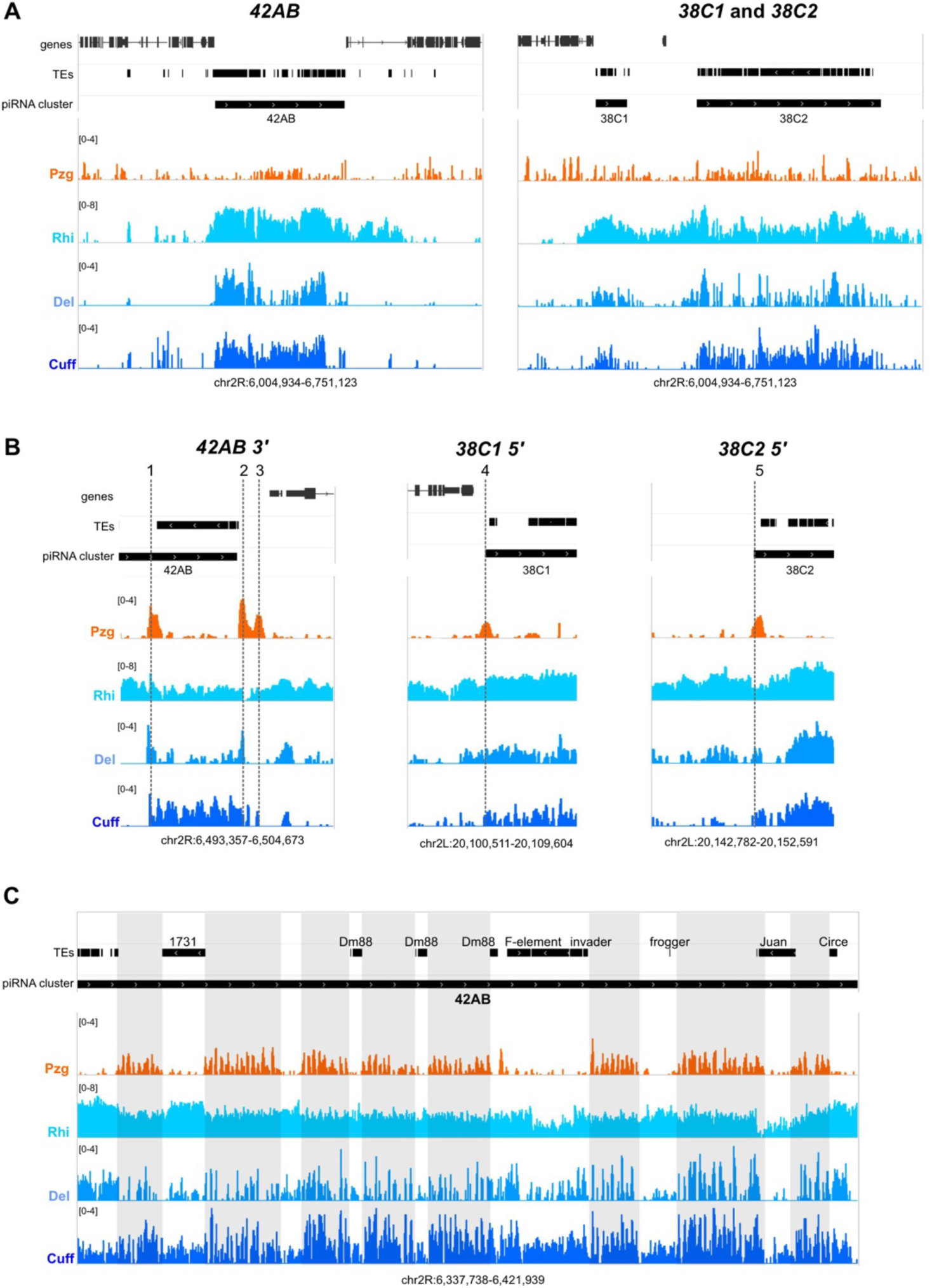
X-Chips on 42AB and 38C1/C2 piRNA clusters. Integrative Genomics Viewer tracks showing the dual-strand piRNA cluster regions *42AB*, *38C1* and *38C2* and ChIP-seq signal (depicted as coverage per million reads) of Pzg (orange) in Pzg-GFP ovaries and Rhi, Del and Cuff (different version of blue color) in control ovaries. ChIP-seq data from the RDC-complex were taken from Baumgartner *et al*., 2024 (Rhi, GEO series GSE202465) and Mohn *et al*., 2014 (Del and Cuff, GEO series GSE55824). Positions are indicated below. **(A)** Overview of entire clusters *42AB* and *38C1/C2*, highlighting specific enrichment of Pzg along the clusters as well as a scattered distribution close by. **(B)** Pzg is highly enriched at cluster borders. Note accumulation at three positions at the 3’ end of cluster 42AB, numbered 1-3, with likewise enrichment of Del notably at peaks 1 and 2. Both clusters *38C1* and *38C2* show strong Pzg enrichment at their 5’ ends, and less prominently of Del and Cuff. **(C)** Close up of cluster region *42AB* depicting TE transcription units. Pzg binding is most predominantly detected in regions without annotated TE transcripts, resembling Del and Cuff.

### Characterization of Pzg-associated proteins in ovaries

With the aim to learn more about Pzg’s role in defending genome stability in the germline, we affinity purified GFP-tagged Pzg protein from ovary lysates using GFP nanobody-coupled beads as well as nanobody-free beads as negative binding control and identified co-associated proteins by mass spectrometry. This seemed reasonable as Pzg has been shown in the past to be part of various protein complexes. Hence, Pzg might influence genome integrity in the germline beyond its role in RDC – Moon/Trf2 mediated transcription initiation. To identify potential interaction partners of Pzg, the immunoprecipitates and their respective negative controls were subjected to differential analysis. Only proteins that were detected in all three replicates (log_2_FC>1 and p_adj_ <0.1) were further considered. In total, we identified 435 significantly enriched proteins, including known Pzg co-factors or proteins expected to interact with Pzg based on earlier analyses (Figure 8A, Table S1). For example, a substantial fraction classified as chromatin factors, e.g. Iswi, CP190, Chro or Heph, which were previously found to interact with Pzg in other tissues (Eggert *et al.,* 2004; Bohla *et al*., 2014; Lowe *et al.,* 2014) (Figure 8A, Table S1). Consistent with the isolation of Pzg interactors from ovaries, a decent fraction has been involved in gametogenesis, i.e. oogenesis, spermatogenesis and related processes. Interestingly, roughly 40% of those were RNA-binding proteins, compared to 14% in general (Table S1). We isolated several components from Notch, JAK/Stat, EcR or Wnt signalling pathways that depend on Pzg activity (Figure 8A, Table S1). Moreover, we also observed a large set of proteins involved in stress response and programmed cell death, in agreements with known Pzg activities (Zimmermann *et al.,* 2015; Kober *et al.,* 2019) (Figure 8A, Table S1). In addition, we co-purified factors involved in post-transcriptional gene silencing, and more precisely, with a well-established role in combatting transposon activity, including Piwi, Hsp83, Tudor-SN, Vasa or Nup358 to name just a few (Figure 8A,B, Table S1). Most interactors in this category are implicated in the Piwi-mediated network of transposon defense (reviewed in Ozata *et al.,* 2019). Piwi is the central player in the piRNA pathway. By recruiting piRNAs, it forms a silencing complex that causes co-transcriptional silencing or post-transcriptional decay of the transposon. To validate this interaction *in vivo*, we precipitated GFP-tagged Pzg protein from ovary extracts and indeed co-precipitated Piwi protein detected by a Piwi specific antibody (Figure 8C). In fact, ovaries from *piwi* mutants are striking phenocopies of those from a *pzg* knockdown: They were described as atrophied and rudimentary, and only contain germaria-like structures. Moreover, these germaria also accumulate a surplus of undifferentiated phospho-Smad positive GSCs in some instances (Jin *et al.,* 2013; Ma *et al*., 2014). However, no direct pairwise protein-protein interactions were detected in a yeast two-hybrid approach (Figure 8D), suggesting that Putzig and Piwi might be partners in Piwi-mediated protein complexes without direct protein-protein contacts.

**Figure 8:**
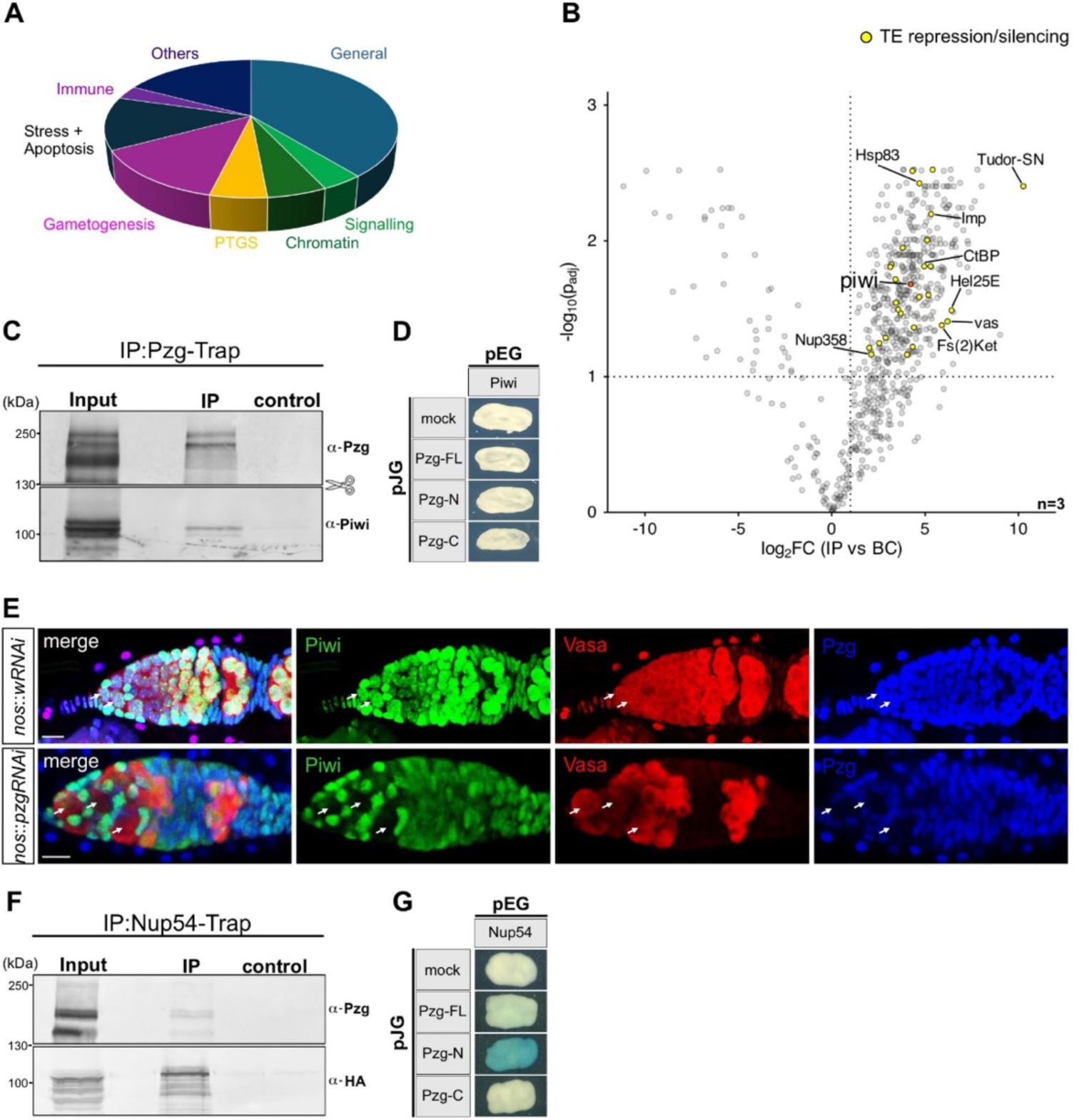
Pzg proteome in ovaries includes Piwi network proteins. (**A**) Pie chart summarizing the proportion of Pzg partners in ovaries belonging to the given categories based on their annotation. For factors falling into several categories, only one is displayed. For details, see Table S1. (**B**) Vulcano Blot showing Pzg-associated proteome in ovaries gained with co-IP mass spectrometry. The dashed lines mark log_2_FC>1 and p_adj_<0.1. Yellow points indicate identified proteins with a known function in transposon repression and/or silencing. Piwi is marked in orange. Results were gained from 3 biological replicates. (**C**) Western Blot analysis of a GFP-tagged Pzg immunoprecipitation experiment from ovaries, demonstrating co-precipitation of Piwi. Input, 10% of IP-trap; control is Trap with agarose beads only. (**D**) Yeast two-hybrid interaction assays revealed no physical interaction between Piwi and Putzig proteins. (**E**) Nuclear Piwi protein is absent in GSC’s upon *pzg* knock down: Co-staining of Piwi (green), Vasa (red) and Pzg (blue) in 0-3 days old germaria using the respective antibodies. In the control (upper row, *nos*-Gal4::UAS-shRNA-*w*) Piwi can be detected in Vasa positive GSCs (arrows), whereas *pzg*-depleted GSCs (lower row, *nos*-Gal4::UAS-shRNA-*pzg*) show a loss of nuclear Piwi (arrows). Scale bars, 10 µm. (**F**) Western Blot of co-immunoprecipitation using mcherry-HA-Nup54 tagged protein from ovarian protein extracts, detected with ⍺-HA and ⍺-Pzg. Input, 10% of IP-trap; binding control were agarose beads. (**G**) Yeast two-hybrid interaction assays revealed an interaction between Nup54 and the N-terminal part of Pzg.

### Nuclear Piwi depends on Pzg

Loading Piwi with piRNA is an essential step for the nuclear entry of Piwi-piRNA complexes (Saito *et al.,* 2010; Yashiro *et al.,* 2018). Our mass spectrometry screen revealed besides Piwi protein itself also various co-purified proteins with an established role in the piRNA export or piRNA biogenesis, like e.g. Vasa, Hsp83, Nup358, or Fs(2)Ket (Figure 8B, Table S1).

Strikingly, >20 of the co-purified proteins have an established role in the nuclear pore complex, including some involved in Piwi-mediated TE silencing like Nup358 (Figure 8B, Table S1) (Handler *et al.,* 2013; Parikh *et al.,* 2018). Hence, we wondered whether Pzg may play a role in Piwi’s nuclear localization. In fact, whereas Piwi and Pzg proteins were co-localized in the nuclei of GSCs of germaria in the control, Piwi was nearly absent in these nuclei upon *pzg* knock down, indicating its importance for Piwi’s nuclear entry or stability (Figure 8E). Moreover, the overexpression of HA-tagged Piwi in *pzg-shRNA* depleted germ cells barely restored nuclear Piwi localization and did not rescue the resultant atrophied *pzg* mutant phenotype, in support of Pzg’s role in Piwi’s nuclear localization (Figure S2).

Several nucleoporins are important for piRNA export and biogenesis (Handler *et al.,* 2013; Parikh *et al.,* 2018; Munafò *et al.,* 2021). For example, Nup54 is required for the export of piRNA-precursors, whereas knockdown of Nup358 in the germline prevents Piwi’s nuclear entry and piRNA biogenesis (Parikh *et al*., 2018). In order to test Pzg’s ability to interact with Nup54, we performed co-IPs with ovary lysates, using flies with a C-terminally labelled Nup54 locus. Not only could we co-precipitate Pzg with Nup54 *in vivo*, moreover we observed a direct interaction of Nup54 with the N-terminal part of Pzg in the yeast two-hybrid approach (Figure 8F,G). For technical reasons, we could not test for direct interaction with Nup358, as the protein was too large for being expressed with the yeast transformation vector. However, the manifold molecular interactions of Pzg with components of the nuclear pore complex suggest some role for Pzg in Piwi-piRNA nuclear localization.

### Neither piRNA amplification via the ping-pong cycle nor subsequent primary piRNA biogenesis are regulated by Pzg

As Pzg function is essential in germ cells for nuclear accumulation of Piwi, we wondered whether Pzg may play a role in piRNA biogenesis and Piwi loading. After their nuclear export, the long piRNA precursors serve as template for the generation of secondary piRNAs in an amplification process known as ‘ping-pong cycle’. This process takes place in the nuage, a perinuclear structure in germ cells, and involves the two Piwi paralogs, Aubergine (Aub) and Argonaute (Ago), both of which bind and trim piRNA transcripts (reviewed in Czech *et al.,* 2018). Even though we had co-purified Aub as well as Vasa, another component driving the ping-pong cycle, in our mass spectrometry screen, we were unable to detect Pzg in co-immunoprecipitates with GFP-tagged Aub and Vasa, respectively (Figure 9A,B). Moreover, both Aub and Ago proteins retained their normal localization in the perinuclear nuage in Pzg depleted germline cells (Figure 9C), indicating that Pzg is not a major player for secondary piRNA generation.

**Figure 9:**
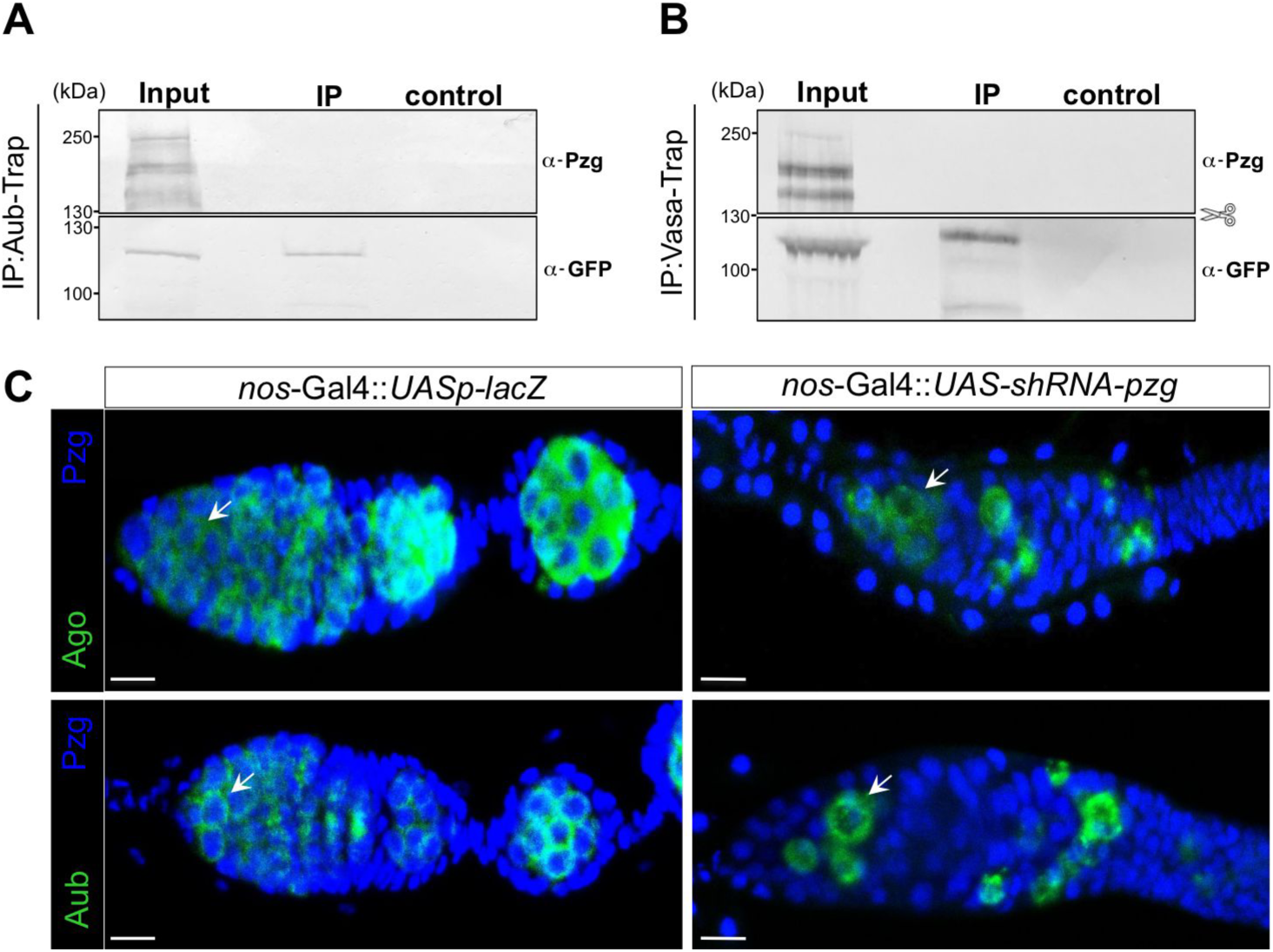
Pzg is not involved in secondary piRNA biogenesis. (**A-B**) Western Blot analysis of GFP-Aub and GFP-Vasa tagged co-immunoprecipitation experiments from ovarian protein extracts. Whereas both tagged proteins could be trapped, no Pzg protein could be co-precipitated. Input, 10% of IP-trap; binding control were agarose beads. (**C**) Confocal images showing Ago and Aub localization in germaria. Both proteins can be detected in the nuage around nuclei of germline cells (green, arrows) in the control (*nos*-Gal4::UASp-*lacZ*) as well as upon knock down of *pzg* in the germline (*nos*-Gal4::UAS-shRNA-*pzg*). Pzg protein is shown in blue. Scale bars, 10 µm.

Eventually, phased piRNA production takes place on the mitochondrial outer membrane, involving the RNA-helicase Armitage (Armi), the endonuclease Zucchini (Zuc), as well as Gasz and Papi, which together split and trim the piRNA intermediates into mature piRNAs loaded onto Piwi (reviewed in Czech *et al.,* 2018; Ozata *et al*., 2019). Albeit none of these factors had shown up in our mass spectrometry assay, we still tested complex formation with Pzg via co-IP assays using ovary lysates from flies expressing the respective tagged protein versions. Indeed, Pzg was co-purified with each of the four components, Armi, Zuc, Gasz and Papi (Figure S3A-D). However, direct protein-protein interactions could not be observed in a yeast two-hybrid assay (Figure S3E). Together, these findings suggest that Pzg is not an integral component of the piRNA-biogenesis machinery, albeit Pzg may be involved in Piwi-piRNA nuclear import from the place of generation.

### Pzg plays a role in TE co-transcriptional gene silencing

Once the Piwi-piRNA complex enters the nucleus and forms a duplex with nascent TE transcripts, the active Piwi complex (Piwi*) with Mael and Arx is formed, to allow the docking of the SFiNX-complex via the binding of Nxf2 (Portell-Montserrat *et al.,* 2025). SFiNX then initiates co-transcriptional gene silencing, by allowing chromatin modifiers Lsd1 and Egg to establish heterochromatin via the docking of HP1a (Sienski *et al*., 2012; Batki *et al*., 2019; Fabry *et al*., 2019; Murano *et al.,* 2019; Zhao *et al.,* 2019; Bence *et al*., 2024). Based on the observation that Pzg associated with Piwi in the ovary (Figure 8B,C), we wondered whether Pzg might be involved in co-transcriptional silencing of TEs. Thus, we next tested whether we can include Pzg molecularly in the Piwi-associated silencing complex by performing co-IPs using tagged proteins from corresponding ovary lysates and yeast two-hybrid assays, respectively. In accordance with the observed association of Pzg and Piwi proteins, we were able to co-precipitate Mael and Pzg from ovary extracts (Figure 10A). Moreover, we found a weak physical interaction of Pzg’s N-terminal domain with Mael in the Yeast two-hybrid assay (Figure 10E), suggesting close contact of Pzg with Piwi*. Moreover, by co-IP we found Nxf2 together with Pzg, but no direct protein-protein interactions (Figure 10B,E), suggesting that Pzg is also present in the Piwi*/SFiNX complex. Finally, the co-IP of both Lsd1 and HP1a together with Pzg strongly supports a role for Pzg in the Piwi-mediated co-transcriptional silencing machinery (Figure 10C,D). Whereas yeast two-hybrid assay did not uncover direct contacts between Pzg and HP1a, we observed a robust molecular interaction of Pzg’s N-terminal domain with Lsd1, supporting the notion that Pzg links the PIWI*/SFiNX complex with Lsd1 to mediate transcriptional silencing (Figure 10E). To further test this idea, we analyzed H3K4me2 levels upon *pzg* knock down in GSCs. Indeed, whereas the control GSCs displayed uniformly moderate H3K4me2 levels, GSCs depleted for *pzg* activity showed a strongly clustered accumulation of nuclear H3K4me2 staining (Figure 10F,G). In sum these data indicate a role for Pzg in the Piwi-mediated co-transcriptional TE silencing: by docking to the Piwi*/SFiNX complex, Pzg may recruit Lsd1 for subsequent demethylation of H3K4 at the active TE promoter, thereby promoting transcriptional silencing. The subsequent H3K9 trimethylation then serves as molecular mark for HP1a binding and the resultant chromatin compaction in consequence. Eventually, enduring transposon silencing occurs via heterochromatin formation of TE loci.

**Figure 10:**
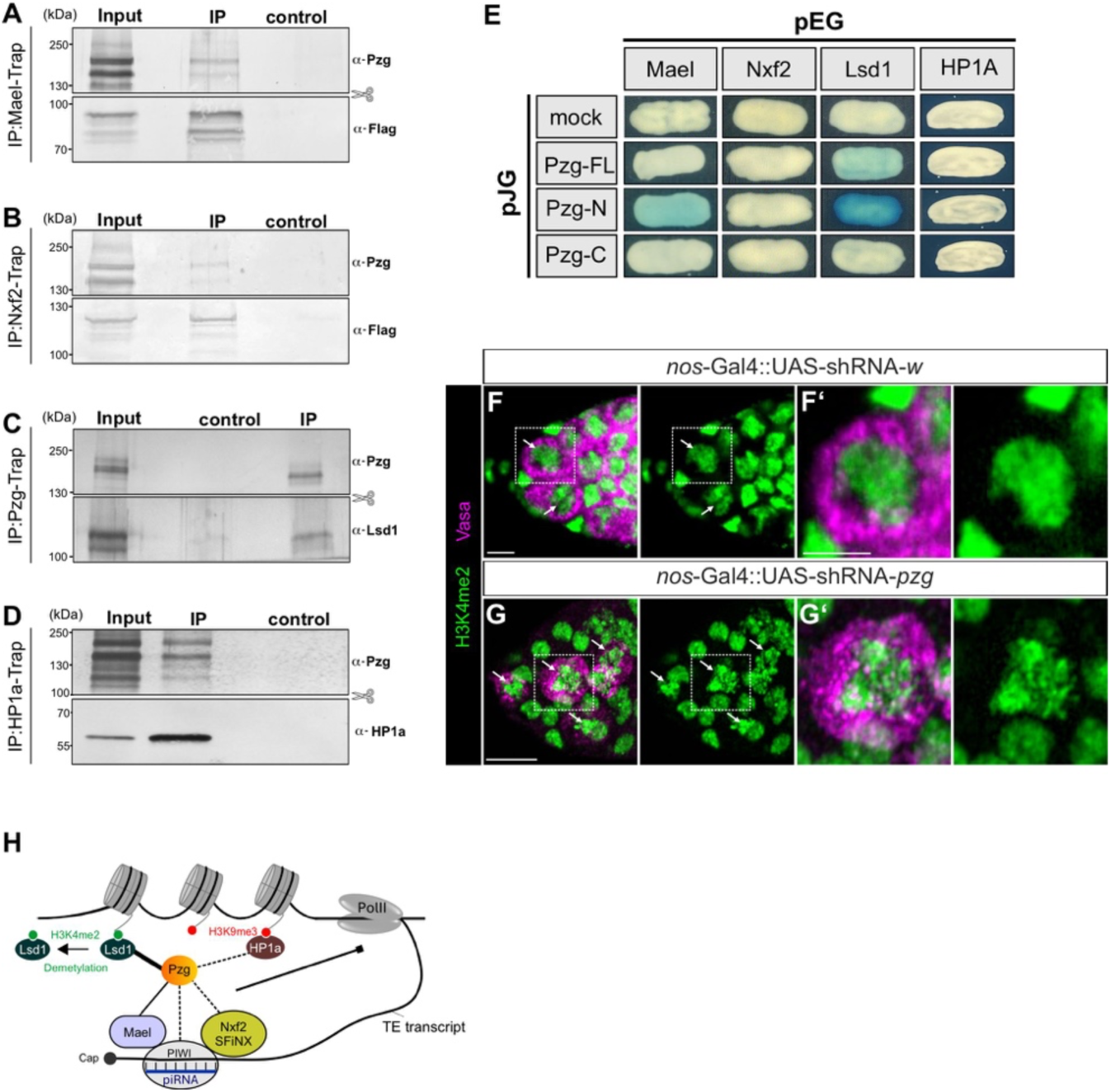
Pzg links Piwi* with Lsd1. (**A-D**) Western Blot analysis of co-immunoprecipitations using tagged proteins as indicated, Mael (A), Nxf2 (B), Pzg (C) and HP1a (D) from ovarian protein extracts, detected with anti-Flag or protein specific antibodies as depicted. Mael, Nxf2 and Pzg were immunoprecipitated from Flag-GFP-tagged lines by GFP-Trap and HP1a from an RFP-tagged HP1a line by RFP-Trap, using nanobodies to capture the tagged proteins. Input, 10% of IP-trap; binding control were agarose beads. (**E**) Yeast two-hybrid tests were performed testing for direct protein-protein interactions. Mael, Nxf2, Lsd1 and HP1a were fused with the DNA binding domain lexA in pEG-vector, whereas Pzg proteins (Pzg-FL, full length), N-terminus (Pzg-N, aa 1-543) and C-terminus (Pzg-C, aa 544-996) were fused to the acidic domain in the pJG vector. Pzg strongly interacts via its N-terminal domain with Lsd1 and also weakly with Mael. (**F-G’**) Detection of H3K4me2 (green) in 0-3 days old germaria with germline cells labelled by α-Vasa (magenta). Compared to the control (*nos*-Gal4::UAS-shRNA-*w*; F-F’), level and distribution of H3K4me2 is changed in *pzg*-depleted GSCs (*nos*-Gal4::UAS-shRNA-*pzg,* G-G’) (arrows in F,G). F’ and G’, enlargements from F and G as indicated by dotted line. Scale bars, 10 µm. (**H**) Schematic of the co-transcriptional silencing complex including confirmed contacts to Pzg.

## Discussion

In this study, we uncover a two-tier activity of Pzg in the process of Piwi-piRNA mediated transposon silencing in *Drosophila*. We show that Pzg, on the one hand, promotes transcription of piRNA clusters by coupling the Trf2/Moon transcription initiation complex to the Rhi/Del/Cuff complex. On the other hand, Pzg is involved in the co-transcriptional silencing of transposon loci by connecting the Piwi*-SFiNX complex with the demethylase Lsd1, thereby allowing promoter inactivation. H3K4me2 removal by Lsd1 at the transcription start site is followed by H3K9me3 deposition via methyltransferase Egg/Wde, which serves as a chromatin mark for HP1a (reviewed in Czech *et al.,* 2018; Di Stefano, 2022). Accordingly, Pzg promotes heterochromatin formation at TE loci. Thus, this work fills important gaps in our current knowledge, as the physical associations amongst the respective components involved have remained unresolved so far.

Pzg acts as a multi-functional protein. As integral component of several multi-protein complexes, it displays a wide range of activities in the regulation of genes and chromatin. Earlier we showed that Pzg is required for germline stem cell (GSC) survival and differentiation in the female germline. Depletion of Pzg activity in the germline resulted in the formation of a new niche-like structure with ectopic GSCs due to the release of morphogens like Dpp as a consequence of cell death (Kober *et al.,* 2019). Several evidence indicate the involvement of Notch, JNK or Jak/Stat signaling pathways in this process (Figures 1-3). Yet, the similarity between *pzg* and *piwi* mutant phenotypes led us to investigate the connection between the two (Jin *et al.,* 2013; Ma *et al*., 2014; Kober *et al.,* 2019). In agreement with a role of Pzg in the Piwi-piRNA pathway, we observed a marked increase in TE activity upon *pzg* knock down with a strong bias for germline specific retrotransposons (Figure 4).

Whereas TE transcription increased in the absence of *pzg*, piRNA precursor transcription decreased in a germline specific manner, notably of the largest dual-strand piRNA cluster *42AB* as well as *80F* (Figure 5). These piRNA clusters bear heterochromatic marks H3K9me3 and H3K27me3, serving as anchors for the HP1 homologue Rhino (Rhi), which facilitates non-canonical transcription of piRNA precursor transcripts. In fact, dual-strand piRNA clusters lack conventional promoters and transcription start sites, respectively, as well as typical transcription termination signals (reviewed in Czech et al., 2018; Ozata *et al.,* 2019). Instead, the Rhino/Deadlock/Cutoff complex (RDC) recruits the germline specific transcription-initiation complex Moon-Trf2. RDC tethers the Moon-Trf2 transcription-initiation complex to both strands of piRNA-clusters to initiate dual-strand cluster transcription from multiple sites (Andersen *et al*., 2017). How Moon-Trf2 is recruited to RDC is still unresolved. Close association between Moon and Del was observed, but no physical interactions (Andersen *et al*., 2017; Riedelbauch *et al*., 2025). In fact, Pzg may act as a link to effectively dock Moon-Trf2 transcription initiation to RDC. During this process, Rhi serves as a molecular hub for effector proteins Del and Pzg, with Del recruiting Cuff and Pzg recruiting Trf2-Moon complexes, altogether allowing efficient piRNA precursor transcription from both DNA strands (Figure 6C). Accordingly, the undulated localization of Pzg in the center of the 42AB cluster sparing defined transposons is in line with positioning transcription complexes rather than with transcription elongation (Figure 7).

Involvement of Pzg in the RDC complex was anticipated, as Pzg has been shown before to be an integral part of the Trf2/Dref complex involved in the transcription of replication-related genes (Hochheimer *et al.,* 2002). The direct physical association of Pzg and Trf2 as well as of Pzg, Rhi and Del confirm a role for Pzg as an adapter between RDC and the Moon-Trf2 transcription-initiation complex (Figure 6). Moon specifically stimulates transcription initiation within piRNA clusters but was shown less relevant at the *38C1/C2* clusters that rely on strong flanking promoters (Andersen *et al*., 2017). Here, Cuff plays an important role: by impeding transcription termination and splicing, Cuff enhances read-through, thereby allowing the formation of more than 15 kb transcripts from adjacent promoters (Mohn *et al*., 2014; Zhang *et al*., 2014; Chen *et al*., 2016; Andersen *et al*., 2017). Pzg is also important in this promoter-driven expression, since the knock down impaired transcription of the *38C1* cluster and notably of the *38C2* plus-strand (Figure 5). Perhaps, Pzg assists Cuff activity, in agreement with its association with the RDC members including Cuff (Figure 6). Accordingly, Pzg protein is enriched at the 5’ ends of *38C1/C2* clusters (Figure 7). The relative increase of the minus-strand transcripts, notably at *38C2*, is reminiscent of the observed strand-bias in plus-strand promoter deletions in the *38C1* cluster (Andersen *et al*., 2017). Apparently, *pzg* knock down resembles a promoter loss at the primary Pol II initiation site, indicating its importance for promoter-driven transcription initiation. Interestingly, Pzg is also involved in transcription of the *20A1* piRNA cluster from a conventional promoter that is Rhino-independent (Mohn *et al.,* 2014), in agreement with its role in canonical promoter-driven transcription, suggesting Trf2 involvement in this context as well (Figure 5).

The mass-spectrometry analysis uncovered an association of Pzg with a multitude of proteins, many of which are involved in the Piwi-piRNA pathway, including Piwi itself. This was confirmed by co-IPs from ovary lysates, albeit direct physical interactions are implausible based on the yeast two-hybrid experiments. However, Pzg supported Piwi’s nuclear localization (Figure 8). We attribute this to the lowered levels of piRNA precursor transcripts, due to Pzg’s role in transcription initiation of piRNA clusters, as this restricts the availability of mature piRNAs and hence the formation of Piwi-piRNA complexes ready for nuclear entry. The loading of piRNA onto Piwi is a prerequisite for nuclear translocation, because conformational changes of Piwi triggered by piRNA loading increases its ability to interact with importins (Saito *et al.,* 2010; Yashiro *et al.,* 2018). Thus, an insufficient level of piRNAs in the Pzg depleted germline might hamper the conformational switch of Piwi needed for the further nuclear import. A more active role of Pzg in the nuclear localization of Piwi-piRNA complexes, however, cannot be excluded based on the manifold interactions of Pzg with proteins of the nuclear pore like Nup358, Nup54 or Fs(2)Ket (Figure 8; Table S1). Moreover, although we have no direct evidence for a role of Pzg within the nuage or in the ping-pong cycle during piRNA amplification, the co-IPs of Pzg with several piRNA processing factors including Armi, Zuc or Hsp83 suggest its presence outside the nucleus (Figure S3; Table S1). Further work is required to unravel the potential activity of Pzg in this context.

Our work, however, clearly uncovered an important role for Pzg in the process of co-transcriptional gene silencing. In the nucleus, the Piwi-piRNA complexes scan for nascent TE transcripts. Upon RNA-duplex formation, the Piwi* complex is built together with Arx and Mael, serving as a platform for Nxf2/Nxt1 binding and subsequent association of Panx to build the SFiNX complex (Murano *et al.,* 2019; Portell-Montserrat *et al.,* 2025). Panx is essential for the recruitment of general heterochromatin effectors Lsd1 and Egg/Wde, licensing co-transcriptional silencing by H3K4me2 removal and H3K9me3 deposition (Sienski *et al.,* 2015; Yu *et al.,* 2015; Batki *et al.,* 2019; Fabry *et al.,* 2019; Lepesant *et al.,* 2020; Murano *et al.,* 2019; Osumi *et al.,* 2019; Portell-Montserrat *et al.,* 2025). Here, Pzg comes into play as it links the Piwi*-SFiNX complex with Lsd1, inducing promoter inactivation, i.e. one of the first steps of TE silencing (Figure 10H). Pzg, on the one hand, contacts Mael in the Piwi* complex, and binds Lsd1 which then can act at the nearby TE promoter, perhaps directed by further contacts of Pzg to promoter-associated factors. This model also explains the observed co-immunoprecipitation of Piwi and Lsd1 (Lepesant *et al.,* 2020). Lsd1 mediates H3K4me2 demethylation at transcription start sites, allowing for deacetylation and subsequent methylation of H3K9, and hence promoter inactivation (Shi *et al.,* 2004; Di Stefano *et al.,* 2007; Lepesant *et al.,* 2020). Interestingly, Panx was recently linked to factors essential for Pol II stalling and termination of TE transcription, i.e. the very first intervention and explaining its ability for gene silencing when tethered to nascent transcripts (Sienski *et al.,* 2015; Yu *et al.,* 2015; Liu *et al*., 2025).

In somatic wing cells, loss of Lsd1 negatively affects cell proliferation and increases DNA damage and cell death, resulting from an upregulation of TE activity (Selmi *et al.,* 2025). The effects on tissue size, cell cycle progression and cell death caused by the loss of Lsd1 are like those induced by depletion of *pzg* in the developing wing (Kugler and Nagel, 2007; Zimmermann *et al.,* 2015). Thus, Pzg might act in combination with Lsd1 in somatic tissues as well. It is tempting to speculate that the two act together in a somatic Piwi-piRNA pathway in the larval fat body which is serving TE-silencing during development (Jones *et al.,* 2016).

The hierarchy of events that lead to TE silencing in the germline, however, are still incompletely resolved, involving several additional factors like Sov, Ova/Su(var)2-1 as well as SUMOylation mediated in part by Su(var)2-10 and by Ubc9 (Jancovics *et al.,* 2018; Batki *et al.,* 2019; Benner *et al.,* 2019; Yang *et al.,* 2019; Ninova *et al*., 2020; Andreev *et al.,* 2022; Portell-Montserrat *et al.,* 2025). Su(var)2-10 directly binds to Piwi promoting SUMOylation of Piwi and of other TE-silencing factors, including the methyltransferase Eggless (Egg) and its co-factor Windei (Wde) (Ninova *et al.,* 2020; Ninova *et al.,* 2023; Bence *et al.,* 2024). Egg/Wde induce trimethylation of H3K9, which is recognized and bound by HP1a to cause heterochromatin formation and enduring chromatin compaction (reviewed in Meyer-Nava *et al.,* 2020). Panx SUMOylation, however, allows the binding of multi-Zinc-finger protein Small ovary (Sov), which enhances heterochromatin formation by supporting the recruitment of HP1a to the chromatin (Jancovics *et al.,* 2018; Andreev *et al*., 2022). Interestingly, Pzg is SUMOylated as well (Ninova *et al.,* 2023), however it is unclear to date whether the SUMOylation plays a role in Pzg-mediated co-transcriptional silencing of TE loci, perhaps by stabilizing protein-protein contacts.

We propose the following sequence of events, some of which presumably occur simultaneously (Figure 11). (1) Pol II-mediated transcription of TEs occurs from active promoters in active chromatin. (2) Heteroduplex formation between TE transcript and piRNA in Piwi-piRNA complex. (3) Formation of Piwi* by recruitment of Arx and Mael. (4) Piwi* binds Nxf2/Nxt1 which recruits Panx. (5) Panx induces stalling of Pol II and termination of TE transcription. (6) Pzg associates with the Piwi* complex by binding to Mael and recruits Lsd1 to the promoter. (7) Lsd1 demethylates H3K4me2/3 at the transcription start site, thereby inhibiting promoter activity and allowing H3K9 methylation. (8) Piwi binds to Su(var)2-10 and SUMOylation of Piwi and other Piwi* complex members occurs. (9) Recruitment of SUMOylated Egg/Wde, which deposit H3K9me3 marks on chromatin that are bound by HP1a. (10) The link between Lsd1 and HP1a by Ova may stabilize the status. (11) SUMOylated Panx binds Sov, which supports the recruitment of HP1a to the chromatin. (12) Enduring heterochromatin compaction by HP1a.

**Figure 11:**
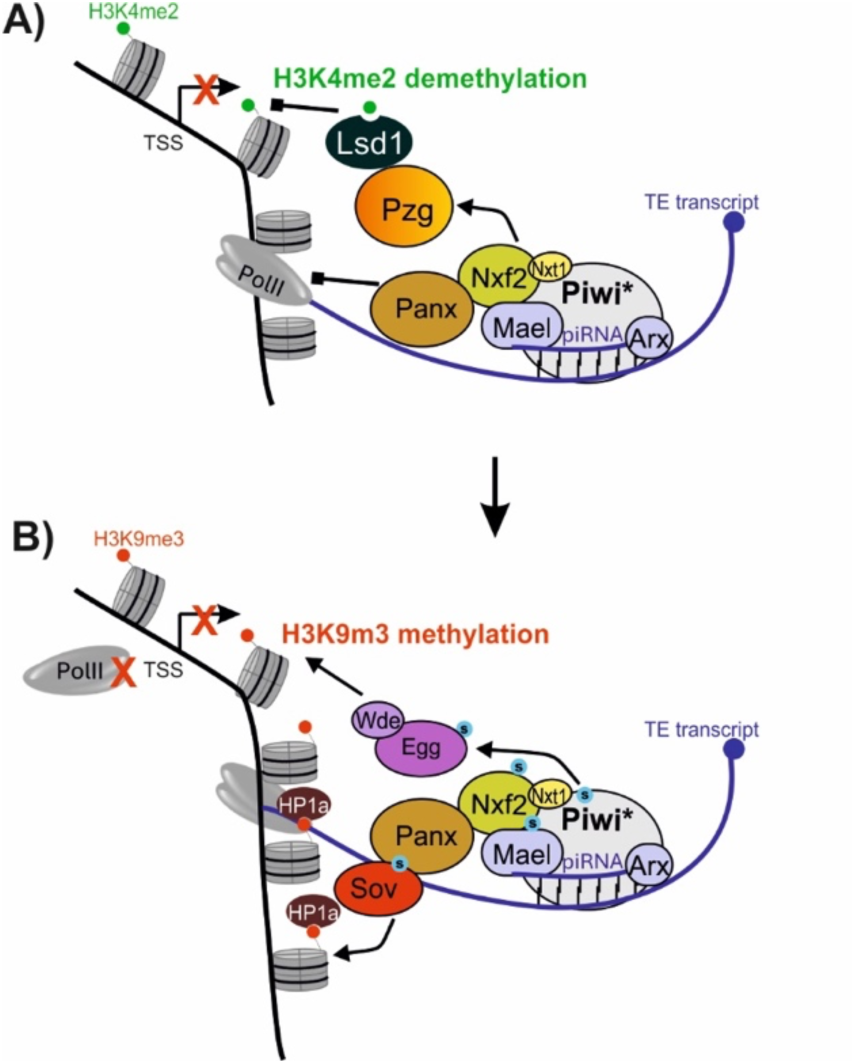
Scheme of proposed sequencing of events occurring at TE loci. **(A)** Initiation of co-transcriptional TE silencing by Panx mediated transcription termination, and Pzg mediated recruitment of Lsd1, inducing promoter silencing. Whilst Pzg associates the Piwi* complex as well as Nxf2, Panx contacts Nxf2. **(B)** Implementation of TE silencing by Egg/Wde induced H3K9me3 methylation, recruiting HP1a to the chromatin supported by Sov, causing enduring heterochromatin formation. SUMOylation (S) promotes Egg/Wde association. For simplicity, not all direct contacts are displayed and several factors were omitted. Further details, see text.

## Supporting information

Key Resources Table

## Acknowledgements

We thank J. Abrams, S. Artavanis-Tsakonas, J. Brennecke, R. Glaser, E. Hafen, L. Johnston, R. Lehmann, P. Rørth for sharing numerous fly stocks and reagents. We obtained fly stocks from the Bloomington stock center (NIH P400D018537) and Vienna *Drosophila* resource Center (VDRC) at Vienna BioCenter Core Facilities (VBCF), member of the Vienna BioCenter (VBC), Austria, as well as antibodies from the Developmental Studies Hybridoma Bank created by NICHD of the NIH and maintained at the University of Iowa (Department of Biology, Iowa City, IA 52242). We thank S. Kugler for the preparation of larval nuclear extracts and L. Schneider for helping with the rescue assays. We are grateful to J. Pfannstiel from the Mass Spectrometry Unit at the Core Facility of the University of Hohenheim for the MS data. We would like to thank the EMBL GeneCore facility, especially V. Benes, J. Landry and M. Boulanger, for their service and expertise. We are indebted to P. Schlüter, K. Feistel and S. Lemke for generous access to equipment and the LSM Zeiss microscope. We are very grateful to A. Preiss for constant support, helpful and stimulating discussions and critical comments on the manuscript.

## Author contributions

Conceptualization: A.C.N.; Data curation: L.H., A.C.N.; Funding acquisition: A.C.N.; Investigation: L.H., L.K., M.Z., S.S., M.E., A.C.N.; Methodology: L.H., L.K., M.Z., S.S.; Supervision: A.C.N.; Validation: L.H., L.K., M.Z., A.C.N.; Visualization: L.H., L.K., M.Z., S.S., A.C.N.; Project administration: A.C.N.; Writing-original draft: A.C.N.; Writing – review and editing: L.H., L.K., M.Z., S.S., M.E., A.C.N.

## Data Availability

The mass spectrometry proteomics data have been deposited to the ProteomeXchange Consortium via the PRIDE partner repository with the following dataset identifiers: PXD075330 and 10.6019/PXD075330. RNA sequencing data have been deposited to GEO under the accession numer GSE324254. Any other data is given in the manuscript, in supplemental files and tables and in Key Resources Table.

