## Supplementary material for "*putzig* safeguards genome integrity by contributing to the Piwi-mediated repression of transposon activity in the female germline of *Drosophila* in a two-tiered fashion": Key Resources Table

| Reagent or Resource | Source | Identifier |
|---|---|---|
| **Antibodies and Dyeing reagents** | | |
| Goat anti-Mouse Alkaline Phosphatase | Jackson ImmunoResearch Laboratories | Cat# 115-055-003; RRID: AB_2338528 (1:1000 WB) |
| Goat anti-Rabbit Alkaline Phosphatase | Jackson ImmunoResearch Laboratories | Cat# 111-055-003; RRID: AB_2337947 (1:1000 WB) |
| Goat anti-Rat Alkaline Phosphatase | Jackson ImmunoResearch Laboratories | Cat# 112-055-003; RRID: AB_2338148 (1:1000 WB) |
| Goat anti-Mouse Cy3 | Jackson Immuno Research | Cat# 115-165-166; RRID: AB_2338692 (1:250 IHC) |
| Goat anti-Rabbit Cy3 | Jackson Immuno Research | Cat# 111-165-144; RRID: AB_2338006 (1:250 IHC) |
| Goat anti-Rat Cy3 | Jackson Immuno Research | Cat# 112-165-167; RRID: AB_2338251 (1:250 IHC) |
| Goat anti-Mouse FITC | Jackson Immuno Research | Cat# 115-095-166; RRID: AB_2338601 (1:250 IHC) |
| Goat anti-Guinea Pig Alexa 647 | Jackson Immuno Research | Cat# 106-605-003; RRID: AB_2337446 (1:250 IHC) |
| Guinea Pig anti-Pzg | own group | Kugler and Nagel, 2007 (1:500 IHC, WB) |
| Mouse anti-Ago | Brennecke lab | Brennecke *et al.*, 2007 (1:50 IHC) |
| Mouse anti-Aub | Brennecke lab | Brennecke *et al.*, 2007 (1:50 IHC) |
| Mouse anti-Flag M2 | Sigma-Aldrich | Cat# F3165; RRID: AB_259529 (1:1000 WB) |
| Mouse anti- $\gamma$H2AV | DSHB | Cat# UNC93-5.2.1: RRID: AB_2618077 (1:2000 IHC) |
| Mouse anti-Hts | DSHB | Cat# 1B1: RRID: AB_528070 (1:20 IHC) |
| Mouse anti-Lamin C | DSHB | Cat# lc28.26, RRID: AB_528339 (1:20 IHC) |
| Mouse anti-Piwi | Brennecke lab, Vienna, Austria | Brennecke *et al.*, 2007 (1:50 IHC) |
| Mouse anti-Piwi | Santa Cruz Technology | Cat#sc-390946; RRID: AB_1125916 (1:200 WB) |
| Normal Goat Serum (NGS) | Jackson ImmunoResearch Laboratories | Cat# 005-000-121; RRID: AB_2336990 |
| Rabbit anti-Di-Methyl-Histone H3 (Lys4) (C64G9) | Cell Signaling Technology | Cat#9725; RRID: AB_10205451 (1:500 IHC) |
| Rabbit anti-GFP | ChromoTek GmbH | Cat# pabg1; RRID: AB_2749857 (1:250 WB) |
| Rabbit anti-GFP XChIP grade | Thermo Fisher Scientific | Cat# A-11122; RRID: AB_221569 (10 $\mu$g per XChIP) |
| Rabbit anti-HA | NEB Biolabs | Cat#3724; RRID: AB_1549585 (1:100 IHC) |
| Rabbit anti-H2A | Glaser lab, New York, USA | Leach *et al.*, 2000 (1:2000 WB) |
| Rabbit anti-$\gamma$H2AX (pSer139) | Trevigen, Inc./Novus | Cat# NB100-384; RRID: AB_10002815 (1:1000 WB) |

| | | |
|---|---|---|
| Rabbit anti-LexA | BioAcademia | Cat#61-001; (1:3000 WB) |
| Rabbit anti-PH3 | Cell Signaling Technology | Cat#9701; RRID: AB_331535 (1:50 IHC) |
| Rabbit anti-phospho-Smad2 (Ser465/467) | Cell Signaling Technology | Cat#3108; RRID: AB_490941 (1:50 IHC) |
| Rat anti-HA clone 3F10 | Roche | Cat#11867423001: RRID: AB_390918 (1:500 WB) |
| Rat anti-Pzg | own group | Kugler and Nagel, 2007 (1:500 WB) |
| Rat anti-Vasa | DSHB | Cat# anti-Vasa; RRID: AB_760351 (1:50 IHC) |
| Vectashield Mounting Medium | Vector Laboratories | Cat# H-1000; RRID: AB_2336789 |
| **Experimental Models: *Drosophila melanogaster* strains** | | |
| **Tagged strains/Reporter/mutant and control** | | |
| *w;;**armi**_EGFP/TM3, Ser* | Vienna Drosophila Resource Center | VDRC (JB-stock) 313238 |
| *w; EGFP_**aub** [attP40];;* | Vienna Drosophila Resource Center | VDRC (JB-stock) 313244 |
| *w;; 3xFLAG/V5/Precission/GFP_**Cuff** [attP2]/TM3, Sb;* | Vienna Drosophila Resource Center | VDRC (JB-stock) 313269 |
| *w;; 3xFLAG/V5/Precission/GFP_**Del** [attP2]/TM3, Sb;* | Vienna Drosophila Resource Center | VDRC (JB-stock) 313271 |
| *w;; GFP_**Gasz** [attP2]/TM3, Sb;* | Vienna Drosophila Resource Center | VDRC (JB-stock) 313277 |
| *w; 3xFlag/GFP_**mael** [attP40]/CyO;;* | Vienna Drosophila Resource Center | VDRC (JB-stock) 313298 |
| *w; ; 3xFLAG/V5/Precission/GFP_CG12721 (**moon**); hs-line [attP2]/TM3,Sb;* | Vienna Drosophila Resource Center | VDRC (JB-stock) 313251 |
| *w; **Nup54** mCherry/3xHA/Precission/4xMYC [attP40]/CyO;;* | Vienna Drosophila Resource Center | VDRC (JB-stock) 313304 |
| *;; FLAG_V5_GFP_**Nxf2** m11-1* | Vienna Drosophila Resource Center | VDRC (JB-stock) 313984 |
| *w; **Papi**_GFP_Precission_V5_3xFLAG [attP40]/CyO;Sb/TM3, Ser;* | Vienna Drosophila Resource Center | VDRC (JB-stock) 313715 |
| *;; FlyFos023014(pRedFlp-Hgr)(**$Z4^{18261}$**::2XTY1-SGFP-V5-preTEV-BLRP-3XFLAG)dFRT = **pzg_GFP*** | Vienna Drosophila Resource Center | VDRC 318536 |
| *w; AID_3xFLAG_V5_Precission_GFP_**rhino**/CyO;;* | Vienna Drosophila Resource Center | VDRC (JB-stock) 313993 |
| *w, moon_FS2Δ28/FM7c; pNASA > LAP **TRF2s**[attP40]/CyO; pNASA > Deadlock-vhhGFP[attP2]/TM3,Ser* | Vienna Drosophila Resource Center | VDRC (JB-stock) 313774 |
| *w; EGFP_**vas** [attP40];;* | Vienna Drosophila Resource Center | VDRC (JB-stock) 313368 |
| *w;; **zuc**_GFP_Precission_V5_3XFLAG [attP2]/TM3,Sb;* | Vienna Drosophila Resource Center | VDRC (JB-stock) 313656 |
| *w;P{NRE-EGFP.S}1* | Bloomington stock center | BL30728; RRID:BDSC_30728 |
| *P{RFP-HP1}3* | Bloomington stock center | BL30562; RRID:BDSC_30562 |
| *P53R-GFPnls* | J. Abrams | Lu *et al.*, 2010 |

| | | |
|---|---|---|
| *nos>NLS_GFP_lacZ_vas-3'UTR_burdock-target [attP2] /TM3,Sb* | Vienna Drosophila Resource Center | VDRC (JB-stock) 313212 |
| *Het-A-lacZ* | | Shpiz *et al.*, 2007 |
| *gypsy-lacZ/CyO* | Vienna Drosophila Resource Center | VDRC (JB-stock) 313222 |
| *pzg*$^{66}$*/TM6B, Ubi-GFP, Tb* | own group | Kugler *et al.*, 2011 |
| *y*$^{1}$*w*$^{67c23}$ | Bloomington stock center | BL6599: RRID:BDSC_6599 |
| **Gal4/UAS lines, hs-line** | | |
| *matα*-Gal4: *w*$^{*}$*; ; P{w*$^{+mC}$*=matalpha4-GAL-VP16}V37;* | Bloomington stock center | BL7063; RRID:BDSC_7063 |
| *nos*-Gal4: *w;; P{w*$^{+mC}$*=GAL4::VP16-nanos.UTR}CG6325*$^{MVD1}$ | Bloomington stock center | BL4937; RRID:BDSC_4937 |
| *bab*-Gal4: *w*$^{*}$*;; P{w*$^{+mW.hs}$*=GawB}bab1*$^{Agal4-5}$*/TM3, Sb*$^{1}$ | Bloomington stock center | BL6802; RRID:BDSC_6802 |
| *c587*-Gal4: *P{w*$^{+mW.hs}$*=GawB}C587, w*$^{*}$*;;* | Bloomington stock center | BL67747; RRID:BDSC_67747 |
| UAS-shRNA-*Dref: y*$^{1}$ *sc*$^{*}$ *v*$^{1}$ *sev*$^{21}$*; P{y*$^{+t7.7}$ *v*$^{+t1.8}$*=TRiP.GL00532}attP2/ TM3, Sb*$^{1}$ | Bloomington stock center | BL36865; RRID:BDSC_36865 |
| UAS-shRNA-*foxo: y*$^{1}$ *sc*$^{*}$ *v*$^{1}$ *sev*$^{21}$*;; P{y*$^{+t7.7}$ *v*$^{+t1.8}$*=TRiP.HMS00422}attP2;* | Bloomington stock center | BL32427; RRID:BDSC_32427 |
| UAS-shRNA-*Nurf301: y*$^{1}$ *sc*$^{*}$ *v*$^{1}$ *sev*$^{21}$*; P{y*$^{+t7.7}$ *v*$^{+t1.8}$*=TRiP.GL00265}attP2* | Bloomington stock center | BL35353; RRID:BDSC_35353 |
| UAS-shRNA-*pzg*: *y*$^{1}$ *sc*$^{*}$ *v*$^{1}$ *sev*$^{21}$*;; P{y*$^{+t7.7}$ *v*$^{+t1.8}$*=TRiP.GL00373}attP2;* | Bloomington stock center | BL35448; RRID:BDSC_35448 |
| *UAS-shRNA-Tom: y*$^{1}$ *sc*$^{*}$ *v*$^{1}$ *sev*$^{21}$*;; P{y*$^{+t7.7}$ *v*$^{+t1.8}$*=TRiP.HMS05421}attP40* | Bloomington stock center | BL66955; RRID:BDSC_66955 |
| *UAS-shRNA-w: y*$^{1}$ *v*$^{1}$*; ; P{y*$^{+t7.7}$ *v*$^{+t1.8}$*=TRiP.HMS00004}attP2/TM3, Sb*$^{1}$ | Bloomington stock center | BL33613; RRID:BDSC_33613 |
| UASp-*lacZ*: *w*$^{+mC,}$ *P{UASp-lacZ}* | P. Rørth | Rørth, 1998 |
| UASp-PiwiHA: *P{UASp-Piwi.HA}/CyO; e vret/TM6b, Tb*$^{1}$ | R.Lehmann | Zamparini *et al.*, 2011 |
| UASt-*Akt1: y*$^{1}$ *w*$^{1118}$*; P{w*$^{+mC}$*=UAS-Akt.Exel}2* | Bloomington stock center | BL8191; RRID:BDSC_8191 |
| UASt-*mir125* | L. Johnston | Caygill and Johnston, 2008 |
| UASt-*Rheb: y*$^{1}$ *w*$^{*}$ *hs-flp;; P{EP(loxP.UAS)y*$^{+}$*.UAS}RhebEP*$^{50.084}$ | E. Hafen | Stocker *et al.*, 2003 |
| UASt-*S6K*$^{TE}$: w; *P{w*$^{+mC}$*=* UAS-*S6K*$^{TE}$*}2* | Bloomington stock center | BL6912; RRID:BDSC_6912 |
| *hs-N*$^{intra}$ | S. Artavanis-Tsakonas | Lieber *et al.*, 1993 |
| **Yeast constructs and vectors** | | |
| pEG202 | E. Golemis | Gyuris *et al.*, 1993 |
| pEG-*armi* (3.6 kb *armi EcoRI/NcoI*) | This work | N/A |
| pEG-*cuff* (1.2 kb *cuff BamHI/XhoI*) | This work | N/A |
| pEG-*del* (3 kb *del EcoRI/XhoI*) | This work | N/A |
| pEG-*gasz* (1.4 kb *gasz EcoRI/XhoI*) | This work | N/A |
| pEG-*Hp1a* (0.6 kb *Hp1a EcoRI/XhoI*) | This work | N/A |
| pEG-*lsd1* (2.7 kb *lsd1 BglII/XhoI*) | This work | N/A |

| | | |
|---|---|---|
| pEG-*mael* (1.4 kb *mael BamHI/XhoI*) | This work | N/A |
| pEG-*moon* (0.5 kb *moon EcoRI/XhoI*) | This work | N/A |
| pEG-*nxf2* (2.6 kb *nxf2 EcoRI/NcoI*) | This work | N/A |
| pEG-*nup54* (1.9 kb *nup54 EcoRI/XhoI*) | This work | N/A |
| pEG-*papi* (1.8 kb *papi EcoRI/NotI*) | This work | N/A |
| pEG-*piwi* (2.6 kb *piwi BglII/NotI*) | This work | N/A |
| pEG-*rhi* (1.3 kb *rhi EcoRI/XhoI*) | This work | N/A |
| pEG-*trf2* (1.9 kb *trf2 BamHI/ XhoI*) | This work | N/A |
| pEG-*zuc* (0.8 kb *zuc NcoI/XhoI*) | This work | N/A |
| pJG4-5 vector | E. Golemis | Vojtek *et al.*, 1993 |
| pJG-Pzg-FL (3kb *Pzg EcoRI/XhoI*) | This work | N/A |
| pJG-*pzg*-N (1.6 kb *pzg EcoRI/XhoI; AS 1-543*) | This work | N/A |
| pJG-*pzg*-C (1.4 kb *pzg EcoRI/XhoI; AS 544-996*) | This work | N/A |
| psH18-34 | E. Golemis | Gyuris *et al.*, 1993 |
| **Critical Commercial kit systems and chemicals** | | |
| Click-iT® EdU Alexa Fluor 488 Imaging Kit for EdU staining | Invitrogen | Cat#C10337 |
| DNA Screen Tape Analysis D1000 Screen Tape and Reagents | Agilent Technologies | Cat # 5067-5582/83 |
| Dynabeads™ Protein G | Thermo Fisher Scientific | Cat# 1003D |
| Binding Control Magnetic Agarose | ChromoTek GmbH | Cat#bmab; RRID: AB_2827548 |
| GFP-Trap® Magnetic Agarose | ChromoTek GmbH | Cat#gtma; RRID: AB_2631358 |
| Myc-Trap® Magnetic Agarose | ChromoTek GmbH | Cat#ytma; RRID: AB_2631370 |
| RFP-Trap® Magnetic Agarose | ChromoTek GmbH | Cat#rtma; RRID: AB_2631363 |
| NextSeq 500/550 Mid-Output v2.5 Kit | Illumina | Cat#20024904/5 |
| Qubit 1X dsDNA HS Assay Kit | Thermo Fisher Scientific | Cat#Q33230 |
| Lamivudine, 3TC | Sigma-Aldrich | Cat#PHR1365 |
| Miracloth filter | Millipore | Cat# 475855 |
| Protease inhibitor cocktail | Roche | Cat# 05892791001 |
| Magnetic Separation Rack | New England Biolabs | Cat# S1506S |
| **Oligonucleotides for cloning of pEG/pJG yeast constructs, 5‘-3‘orientation** | | |
| Armi_UP_*EcoRI:* ATTTGAATTCATGTTCACATACGTTAGC AAG | Microsynth | N/A |
| Armi_LP_*NcoI:* TCATCCATGGTTAGTTCAAATC ATCTGTAGTA | Microsynth | N/A |
| Boot_UP_*BamHI:* CGAGGGATCCCGATGACG GATTTCAATGCATC | Microsynth | N/A |
| Boot_LP_*NcoI:* TGTCCCATGGCTAGTTAAA ACGTTGCAGTTGG | Microsynth | N/A |
| Cuff_UP_Bam*HI:* CGTTGGATCCAGATGAAT TCTAATTACACAATAT | Microsynth | N/A |

| | | |
|---|---|---|
| Cuff_LP_Xho*I:* ATTGCTCGAGTTAAACTATAGAAGACA TGGTT | Microsynth | N/A |
| Del_UP_*EcoRI:* TGGCGAATTCATGGAAAAGTTGGACA | Microsynth | N/A |
| Del_LP_Xho*I:* TTTCCTCGAGTTAATCAAAATTATGTAT ATTG | Microsynth | N/A |
| Gasz_UP_*EcoRI:* AAGTGAATTCATGATGAGCAACCTTTG | Microsynth | N/A |
| Gasz_LP_Xho*I:* TGAGCTCGAGCTATGAGAACCACTTGC | Microsynth | N/A |
| HP1a_UP_*EcoRI*: CATAGAATTCATGGGCAAGAAAATCG ACAACC | Microsynth | N/A |
| HP1a_LP_*XhoI*: GTACCTCGAGTTAATCTTCATTATCAGA GTACCAGGA | Microsynth | N/A |
| Lsd1_UP_*BglII:* CATAAGATCTACATGAAACCCACCCAG TT | Microsynth | N/A |
| Lsd1_LP_Xho*I:* AAATCTCGAGTTACTGTAGCTCCGTAG | Microsynth | N/A |
| Mael_UP_*BamHI:* ACGCGGATCCAGATGGCTCCTAAGAA GCATAG | Microsynth | N/A |
| Mael_LP_ Xho*I :* CCAACTCGAGTTATTTTTTAAGTTTCCC ATCA | Microsynth | N/A |
| Moon_UP_ *EcoRI:* TGGCGAATTCATGGCCAAGATATTGCC | Microsynth | N/A |
| Moon_LP_ Xho*I:* GCTGCTCGAGCTACATTTCCTGGCCTG | Microsynth | N/A |
| Nup54_UP_ *EcoRI:* ATACGAATTCATGTCGTTCTTCGGATC CAACA | Microsynth | N/A |
| Nup54_LP_ Xho*I:* TCCACTCGAGTCACGATTGTCGCAGCT CGGGC | Microsynth | N/A |
| Papi_UP_EcoR*I:* GTTAGAATTCATGTTGCGCAACACGCC TTTCG | Microsynth | N/A |
| Papi_LP_Not*I :* CAATGCGGCCGCCTAATGCGCGCTAGC ACCATTT | Microsynth | N/A |
| Piwi_UP_Bgl*II:* CAACAGATCTTAATGGCTGATGATCAG GGACG | Microsynth | N/A |
| Piwi_LP_Not*I:* AATTGCGGCCGCTTATAGATAATAAAA CTTCTTTTCG | Microsynth | N/A |
| Pzg_UP_*EcoR*I: ACACGAATTCATGAACAACCAACTGAA CCCG | Microsynth | N/A |
| Pzg_LP_AS220_*XhoI*: TAATCTCGAGTTAGGTTCCTGGGGCGA TCTTGGAC | Microsynth | N/A |

| | | |
|---|---|---|
| Pzg_UP_AS221_*EcoRI*: ATTAGAATTCACCCAGAGCAGCGTCCAAGGCA | Microsynth | N/A |
| Pzg_1967_LP_*XhoI*: TATTCTCGAGTTACTCTCCGGGCTCATCGCTTA | Microsynth | N/A |
| Pzg_1970_UP_*EcoRI*: TTTAGAATTCGCAACTGGTAAGGCCGGTGAC | Microsynth | N/A |
| Pzg_LP_*XhoI*: CGGTCTCGAGCTAGTCGGTCTTTGTCTCTGTAAACT | Microsynth | N/A |
| Rhi_UP_EcoR*I:* CGGAGAATTCATGTCTCGCAACCATCA | Microsynth | N/A |
| Rhi_LP_Xho*I:* CTAGCTCGAGTTACTTGGGCACAATGATCCTCA | Microsynth | N/A |
| Trf2_RH_UP_*BamHI*: TCATGGATCCTTATGCAAAACGATATGGTCAGCATAC | Microsynth | N/A |
| Trf2_RH_LP_*XhoI*: GCTACTCGAGTTAGAAGGGCATATCCAATTCGTTG | Microsynth | N/A |
| Zuc_UP_*NcoI*: AATTCCATGGATGTTGATTACCCAAATAATT | Microsynth | N/A |
| Zuc_LP_Xho*I*: CAATCTCGAGCTACTTGAGCTGGATTT | Microsynth | N/A |
| **Oligonucleotides: genotyping** | | |
| Akt1_internally: CCTTTCGATTACCGTGGTCCATTG | Microsynth | N/A |
| eGFP_UP: GGTACGCGCTAGAGTCGAGAG | Microsynth | N/A |
| eGFP_LP: GTGGTATGGCTGATTATGATCTAGAG | Microsynth | N/A |
| Gal4_LP: AAGATGTAGGGCTGTCACCAA | Microsynth | N/A |
| Gal4VP16_UP: GCTGACTAGGGCACATCTGACA | Microsynth | N/A |
| Gal4VP16_LP: CATATCCAGAGCGCCGTAGG | Microsynth | N/A |
| GFP_UP: GCCACAAGTTCAGCGTGTCCG | Microsynth | N/A |
| GFP_LP: TCACCTTGATGCCGTTCTTCTGC | Microsynth | N/A |
| lacZ_UP: CAACGTCGTGAGTGGGAAAACC | Microsynth | N/A |
| lacZ_LP: CCCGTTGCACCACAGATGAAA | Microsynth | N/A |
| Notchintra_UP: CGCCCGTTTTATCCATTGTG | Microsynth | N/A |
| Hsp70_LP: ACGGAGGGACAATTCAATTCAA | Microsynth | N/A |
| pUASt_5‘: CGCACAGTCACGTTATTG | Microsynth | N/A |
| P-end_5‘: CGACGGGACCACCTTATG | Microsynth | N/A |
| pValium20_UP: ACCAGCAACCATCAAC | Microsynth | N/A |

| | | |
|---|---|---|
| pValium20_LP: TAATCGTGTGTGATGCCTACC | Microsynth | N/A |
| pValium22_UP: GGTGATAGAGCCTGAACCAG | Microsynth | N/A |
| pValium22_LP: TAATCGTGTGTGATGCCTACC | Microsynth | N/A |
| Rheb_internally_LP‘: TTCTCCTGGGGATTGCCGTTCT | Microsynth | N/A |
| **Stellaris™ FISH probes, all marked with Quasar® 570** | | |
| **38C2 set (5‘-3‘ orientation)** | | |
| Cluster5_1 TATTTTAATGTGCGCGGTACG | LGC Technologies, Inc. (Hoddesdon, UK) | N/A |
| Cluster5_2 GTCGCTTAATTCAACTTTTCC | LGC Technologies, Inc. (Hoddesdon, UK) | N/A |
| Cluster5_3 CTTCAGTTAGTTCTATTCCTA | LGC Technologies, Inc. (Hoddesdon, UK) | N/A |
| Cluster5_4 TCTTATCTCTTTCTCATTCAT | LGC Technologies, Inc. (Hoddesdon, UK) | N/A |
| Cluster5_5 ATCGCATCCTTTGTATTATTT | LGC Technologies, Inc. (Hoddesdon, UK) | N/A |
| Cluster5_6 CGCACGCGCTTAAATATTTAA | LGC Technologies, Inc. (Hoddesdon, UK) | N/A |
| Cluster5_7 GCTTGGTCGAGTGTGACC | LGC Technologies, Inc. (Hoddesdon, UK) | N/A |
| Cluster5_8 CCACACTAATACAGGCAAATG | LGC Technologies, Inc. (Hoddesdon, UK) | N/A |
| Cluster5_9 GACTGGGATTAGTGTGCC | LGC Technologies, Inc. (Hoddesdon, UK) | N/A |
| Cluster5_10 CGCGCGCTTTATATAGGC | LGC Technologies, Inc. (Hoddesdon, UK) | N/A |
| Cluster5_11 AAATACCGACTGGCTAATGCA | LGC Technologies, Inc. (Hoddesdon, UK) | N/A |
| Cluster5_12 ATATGACTTCGCTGCTGACTA | LGC Technologies, Inc. (Hoddesdon, UK) | N/A |
| Cluster5_13 CAATAGACTCGGATTTCAGGA | LGC Technologies, Inc. (Hoddesdon, UK) | N/A |
| Cluster5_14 CTCCATCAGAGCATTTTATAC | LGC Technologies, Inc. (Hoddesdon, UK) | N/A |
| Cluster5_15 ATCATTTTCCAAATTCGTGCA | LGC Technologies, Inc. (Hoddesdon, UK) | N/A |

| | | |
|---|---|---|
| Cluster5_16 GGCCAATAAAGTTCGATTCGA | LGC Technologies, Inc. (Hoddesdon, UK) | N/A |
| Cluster5_17 GCGAGTCAGATGATGATGTTA | LGC Technologies, Inc. (Hoddesdon, UK) | N/A |
| Cluster5_18 GGTACTTGCAGAAATCCT | LGC Technologies, Inc. (Hoddesdon, UK) | N/A |
| Cluster5_19 CCTCCTGAAAATATATTTGCG | LGC Technologies, Inc. (Hoddesdon, UK) | N/A |
| Cluster5_20 AACTAGTAGAGTTAATCCCTG | LGC Technologies, Inc. (Hoddesdon, UK) | N/A |
| Cluster5_21 AGGATCACACAGACTTTTTCT | LGC Technologies, Inc. (Hoddesdon, UK) | N/A |
| Cluster5_22 GCAAAAGTGGAACAGCATCAC | LGC Technologies, Inc. (Hoddesdon, UK) | N/A |
| Cluster5_23 TCCACTTCGACATATTCTTAA | LGC Technologies, Inc. (Hoddesdon, UK) | N/A |
| Cluster5_24 GGGCACAAATACAGGATGGAA | LGC Technologies, Inc. (Hoddesdon, UK) | N/A |
| Cluster5_25 AGTTAGGTTGGAATCCAAGAC | LGC Technologies, Inc. (Hoddesdon, UK) | N/A |
| Cluster5_26 CAGTGCCATAAAAACCCGAAA | LGC Technologies, Inc. (Hoddesdon, UK) | N/A |
| Cluster5_27 AAATGCTGTAAGATCACCTCC | LGC Technologies, Inc. (Hoddesdon, UK) | N/A |
| Cluster5_28 AGTTGCAATAATGCCGATATT | LGC Technologies, Inc. (Hoddesdon, UK) | N/A |
| Cluster5_29 CCAGTTGATATTGGGGTTAAA | LGC Technologies, Inc. (Hoddesdon, UK) | N/A |
| Cluster5_30 TTTATCACTGTTTTTATGCCA | LGC Technologies, Inc. (Hoddesdon, UK) | N/A |
| Cluster5_31 GATGGGGGGAAAAATGCCAAT | LGC Technologies, Inc. (Hoddesdon, UK) | N/A |
| Cluster *42AB* probes | LGC Technologies, Inc. (Hoddesdon, UK) | Andersen *et al.*, 2017 |
| **Software, Algorithms and Imaging systems** | | |
| Axioskop 2 Plus coupled with Canon EOS 700D camera | Carl Zeiss/Canon | N/A |
| Axiophot stereomicroscope coupled with Pixera PVC 100C camera | Carl Zeiss/Pixera Corporation | N/A |

| | | |
|---|---|---|
| MRC1024 Laser scanning confocal imaging system and Axiophot microscope | Bio-Rad/Zeiss | N/A |
| LSM700 confocal microscope | Carl Zeiss Microscopy | N/A |
| 3i W1 Spinning Disk microscope | 3i Intelligent Imaging Innovations GmbH | N/A |
| SlideBook 6 software for 3i W1 Spinning Disk microscope | 3i Intelligent Imaging Innovations GmbH | N/A |
| LaserSharp 2000 software for MRC1024 | Bio-Rad | N/A |
| Creation of images using LSM700 confocal microscope | ZEN<br>Version SP7 | Carl Zeiss Microscopy GmbH |
| Figure assembly | CorelDRAW®<br>Graphics Suits<br>Version 9 | RRID:SCR_014235 |
| Picture processing and analyzing | Corel Photo Paint®<br>Version 9.337 | N/A |
| Data processing and analyzing | Fiji (Image J)<br>Version ImageJ2 (2.16.0) | RRID:SCR_003070 |
| Filtering ChIP-Seq data | BBMap<br>Version 39.19 | Bushnell B. (sourceforge.net/projects/bbmap/) |
| Intersection with blacklist regions | Bedtools<br>Version 2.31.0 | Quinlan, 2014 |
| ChIP-seq Alignment | Bowtie2<br>Version 2.5.4 | Langmead and Salzberg, 2012 |
| Use of various tools for ChIP-seq data processing and analysis | deepTools<br>Version 3.5.6 | Ramírez *et al.*, 2016 |
| ChIP-seq Quality analysis | FastQC<br>Version 0.12.1 | Andrew S. (www.bioinformatics.babraham.ac.uk/projects/fastqc/) |
| Visualizing of ChIP-seq alignments | IGV<br>Version 2.18.4 | Robinson *et al.*, 2011 |
| Removal of ChIP-seq duplicates | Picard<br>Version 3.3.0 | Broad Institute, 2019. Picard Toolkit. Broad Inst. GitHub Repos |
| Use of various tools for ChIP-seq data processing | SAMtools<br>Version 1.21 | Danecek *et al.*, 2021 |
| Creation of ChIP-seq fragmentation profiles | TapeStation Analysis Software Version 4.1.1 | Agilent Technologies |
| ChIP-seq quality analysis and data processing | Trim Galore<br>Version 0.6.10 | Krueger F. (www.bioinformatics.babraham.ac.uk/ Projects/trim:galore/) |
| Data analysis and visualization | R<br>Version 4.5.1 | R Core Team, 2024 (https://www.R-project.org/) |
| User interface for R | R-studio<br>Version 2025.05.0+496 | Posit team, 2025 (http://www.posit.co/) |
| Creation of diagrams in R | dplyr<br>Version 1.1.4<br>ggplot2 | Wickham *et al.*, 2025: dplyr: https://dplyr.tidyverse.org |

| | | |
|---|---|---|
| | Version 3.5.2<br>tidyverse<br>Version 2.0.0 | Wickham, 2016<br><br>Wickham *et al.*, 2019 |
| IP-MS data analysis | MaxQuant<br>Version 2.6.7.0 | Cox and Mann, 2008 |
| Statistical evaluation of IP-MS data | Perseus<br>2.0.11 | Tyanova *et al.*, 2016 |
| GO analyses of IP-MS data | ShinyGO<br>Version 0.81 | Ge *et al.*, 2020 |
| Workbench version v2.0.7 | bms/Biozym | Cat # 68MIC-HRM |
| **Deposited data** | | |
| Mass Spectrometry datasets shown in the paper | This study | PRIDE: PXD075330 and 10.6019/PXD075330 |
| Pzg-GFP ChIP-seq data shown in this paper | This study | GEO series GSE324254 |
| Rhi ChIP-seq data | Baumgartner *et al.*, 2024 | GEO series GSE202465 |
| Del ChIP-seq data | Mohn *et al.*, 2014 | GEO series GSE55824 |
| Cuff ChIP-seq data | Mohn *et al.*, 2014 | GEO series GSE55824 |
| **Materials and equipment for qRT-PCR** | | |
| Blue S'Green qPCR Kit (for qPCR in *Drosophila* tissue) | Biozym | Cat # 331416 |
| DNaseI | New England Biolabs | Cat # M0303 |
| Dynabeads™ mRNA DIRECT™ Micro Purification Kit | Invitrogen, Thermo Fisher Scientific | Cat # 61021 |
| miRNeasy Micro Kit (for isolation of total RNA from ovaries) | Qiagen | Cat # 217084 |
| RNase-free DNase set | Qiagen | Cat # 79254 |
| ProtoScript II First strand cDNA synthesis Kit | New England Biolabs | Cat # E6560S |
| qScriber™ cDNA Synthesis Kit | highQu | Cat # RTK0104 |
| RNA*later* Stabilization Solution | Thermo Fisher Scientific | Cat # AM7020 |
| Mic Magnetic Induction Cycler | bio molecular systems/Biozym | Cat # 68MIC-2 |
| **Oligonucleotides used for transposon, controls and *tom* qRT-PCR** | | |
| Blood_UP: AACAATAGAAAGAAGCCACCGAAC | Microsynth | Handler *et al.*, 2011 |
| Blood_LP: AGTCATGGACTATTGAGGGTGTTG | Microsynth | Handler *et al.*, 2011 |
| Burdock_UP: TTGTGCAACCAATCGGCTCTC | Microsynth | N/A |
| Burdock_LP: ATGAAGTGGAGGTGCGTTCGC | Microsynth | N/A |
| Copia_UP: CGACAGTGTGGAGGTTGTGCC | Microsynth | Shpiz *et al.*, 2011 |
| Copia_LP: CTTGGAGACGCTTTACGGACAT | Microsynth | Shpiz *et al.*, 2011 |
| CPSF_UP: GGGTTTTCTGGTGACGGGACTTG | Microsynth | N/A |
| CPSF_LP: CGACAATGTGCGATACTCCTCCTG | Microsynth | N/A |
| cyp33_UP: CTCTGCGGACGCACAATTC | Microsynth | PP14577 DRSC FlyPrimerBank |

| | | |
|---|---|---|
| cyp33_LP: TGCAACCAGTCGTCATCTGC | Microsynth | PP14577 DRSC FlyPrimerBank |
| dlp_UP: ATGCGGGTAGTGGAGCTGGTTCT | Microsynth | N/A |
| dlp_LP: GTCCGGCTTATGGCTGGGTTC | Microsynth | N/A |
| F-element_UP: TTGTTGAACAGCATACCACTCC | Microsynth | Czech *et al.*, 2008 |
| F-element_LP: CCAGAGTTGATGAGCCAGTGTA | Microsynth | Czech *et al.*, 2008 |
| gypsy_UP: CCATACCCAGCTATACAGGTGAGATGG | Microsynth | N/A |
| gypsy_LP: GACCGTTTTGATATTGGGGAACAG | Microsynth | N/A |
| HetA_UP: CCAGGCAAGCGGACAAACGA | Microsynth | Shpiz *et al.*, 2011 |
| HetA_LP: GGAGTGATGAGCGGCGGAAA | Microsynth | Shpiz *et al.*, 2011 |
| Hobo_UP: AAT CGG CGC TAT CTA TGG GGA A | Microsynth | N/A |
| Hobo_LP: CACTTGCTCTTCCGCTATCCACAG | Microsynth | N/A |
| Hopper_UP: CAAGATCCCGCAGGCAGCAC | Microsynth | N/A |
| Hopper_LP: TGAGTGGCCCAAGACAGCAAGC | Microsynth | N/A |
| Idefix_UP: ATTCCACAAGACCGTAAACGA AAAG | Microsynth | N/A |
| Idefix_LP: CATATAGGAAAATGTCTGCTGGAAATC | Microsynth | N/A |
| I-element_UP: TGAAATACGGCATACTGCCCCCA | Microsynth | Teo *et al.*, 2018 |
| I-element_LP: GCTGATAGGGAGTCGGAGCAGATA | Microsynth | Teo *et al.*, 2018 |
| jockey_UP: TATTGGCACTGCTCAGGCTAAA | Microsynth | N/A |
| jockey_LP: TTTCTTCGAGCAAGGGCCTTAT | Microsynth | N/A |
| LamC_UP: CTGGAGAACCTCTTGGACACGGA | Microsynth | N/A |
| LamC_LP: CCTCACCGCACAGCAGTTTGTC | Microsynth | N/A |
| mdg1_UP: ATATGTGCGACAAATTTCGTCCAC | Microsynth | Batki *et al.*, 2019 |
| mdg1_LP: CAAGTATCTTTGTCTTGATTACCTCAAC | Microsynth | Batki *et al.*, 2019 |
| R2_UP: ATGCTCCCGAAACAACAAAC | Microsynth | Teo *et al.*, 2018 |
| R2_LP: GCACTGCAGACTTGGTTCAA | Microsynth | Teo *et al.*, 2018 |
| roo_UP: CGTCTGCAATGTACTGGCTCT | Microsynth | Iwasaki *et al.*, 2021 |
| roo_LP: CGGCACTCCACTAACTTCTCC | Microsynth | Iwasaki *et al.*, 2021 |
| SdhA_UP: GGACGGTGTGAACAAGATGAAGGAG | Microsynth | N/A |
| SdhA_LP: CGCTCACAATAGTCATCTGGGCAT | Microsynth | N/A |
| S-element_UP: | Microsynth | N/A |

| | | |
|---|---|---|
| CTGTGCGCCAAGTCATCCTACG | | |
| S-element_LP: CAACGGTTTGCGCCATACTCTG | Microsynth | N/A |
| TART-A1_UP: AATGAACTTTGTCTGCCCTCCCA | Microsynth | Shpiz *et al.*, 2011 |
| TART-A1_LP: ATCTGTCTACTGTCCGCCTTCGCTA | Microsynth | Shpiz *et al.*, 2011 |
| tbp_UP: TAAGCCCCAACTTCTCGATTCC | Microsynth | PP1556 DRSC FlyPrimerBank |
| tbp_LP: GCCAAGAGACCTGATCCC | Microsynth | PP1556 DRSC FlyPrimerBank |
| Tom_UP: ACTGGTCAAACCCCTTCTGC | Microsynth | PD44529 DRSC FlyPrimerBank |
| Tom_LP: ATCTTTGCGCATGTCCTCCA | Microsynth | PD44529 DRSC FlyPrimerBank |
| Transib2_UP: AAACTGCCCGAGTGTAACAGACAAC | Microsynth | N/A |
| Transib2_LP: TTTCGGCGTCCCATTTGCTG | Microsynth | N/A |
| ZAM_UP: CTACGAAATGGCAAGATTAATTCCACT TCC | Microsynth | N/A |
| ZAM_LP: CCCGTTTCCTTTATGTCGCAGTAGCT | Microsynth | N/A |
| 297_UP: AACCCGGAACATTGCCGAGA | Microsynth | N/A |
| 297_LP: AAGCTCCGTCTCTGTCTGCC | Microsynth | N/A |
| 412_UP: AAAGTACGGTCCAATGAAGACG | Microsynth | Czech *et al.*, 2008 |
| 412_LP: GTGGTGATGAGCTGTTGATGTT | Microsynth | Czech *et al.*, 2008 |
| **Oligonucleotides used for piRNA/controls reverse transcription** | | |
| clFlam_RT_plus: GGGCAATAGTACGCTAGTCCG | Microsynth | N/A |
| cl20A1_RT_plus: ATGGGCGATACATTGTGTCAA | Microsynth | N/A |
| cl38C1_RT_plus: CCT CCT CTT CGG ACT CCT ACG | Microsynth | N/A |
| cl38C1_RT_minus: CGT CTT TGC CAA GCA CTC C | Microsynth | N/A |
| cl38C2_RT_plus: CATTTTTCGGCGTCCACAGT | Microsynth | N/A |
| cl38C2_RT_minus: GTGGGCTGTGATAGCTGATAGTGT | Microsynth | N/A |
| cl42AB_RT_plus: CGAAGCCTTAGATCTCGCTCC | Microsynth | Klattenhoff *et al.*, 2009 |
| cl42AB_RT_minus: ACA TCA GGA ACA CAG CGA GGT G | Microsynth | Klattenhoff *et al.*, 2009 |
| cl80F_RT_plus: TCTCCCTGACCAACACGAAAC | Microsynth | N/A |
| cl80F_RT_minus: GCAATCTCGCTGGGAGTGAT | Microsynth | N/A |
| CPSF_RT_1: ATG GCA CTG CAC CCG AAA C | Microsynth | N/A |
| cyp33_RT_1: CAC AAT TGC CGA AAG TTC TCC | Microsynth | N/A |
| sdha_RT_1: TGC TTG CGC CAG TGC TGA | Microsynth | N/A |

| | | |
|---|---|---|
| tbp_1_RT_1:<br>CCC ACA CTC CGC TCA CTC A | Microsynth | N/A |
| dlp_RT_1:<br>GCC TGC ATC CGT GTC CTC | Microsynth | N/A |
| LamC_RT_1:<br>GAT TCC AGA ATG CCA GGT GTA GC | Microsynth | N/A |
| **Oligonucleotides used for piRNA qRT-PCR** | | |
| clFlam_UP:<br>CTGAGGAATGAATCGCTTTGAAGTA | Microsynth | N/A |
| clFlam_LP:<br>GTAGTATTGCCTACTTTTTAGCCGA | Microsynth | N/A |
| cl20A_UP:<br>GTGTAGCCGGCAAAATGCAGC | Microsynth | N/A |
| cl20A_LP:<br>CCCTCAATATGTAGAGTAGTGCGAG | Microsynth | N/A |
| cl38C1_UP:<br>GGT ATG AAG ATT GGA GGC GGC TTG | Microsynth | N/A |
| cl38C1_LP:<br>CCA CCC ACC ACC AAG CTG CTC TA | Microsynth | N/A |
| cl38C2_UP:<br>CTTGGTAACAAGTACCGTATGGTCG | Microsynth | N/A |
| cl38C2_LP:<br>GGACCTTCCCTGCTCCCTACAA | Microsynth | N/A |
| cl42AB_UP:<br>GCGT CCC AGC CTA CCT AGT CAG AA | Microsynth | N/A |
| cl42AB_LP:<br>ACT TCC CGG TGA AGA CTC CTT CC | Microsynth | N/A |
| cl80F_UP:<br>TCGTGAGTCGCGTCAACATCCT | Microsynth | N/A |
| cl80F_LP:<br>GAGGCAGTACCTTTCCCGCTCA | Microsynth | N/A |
