## Supplemental Figures S1-S4 for "*putzig* safeguards genome integrity by contributing to the Piwi-mediated repression of transposon activity in the female germline of *Drosophila* in a two-tiered fashion"

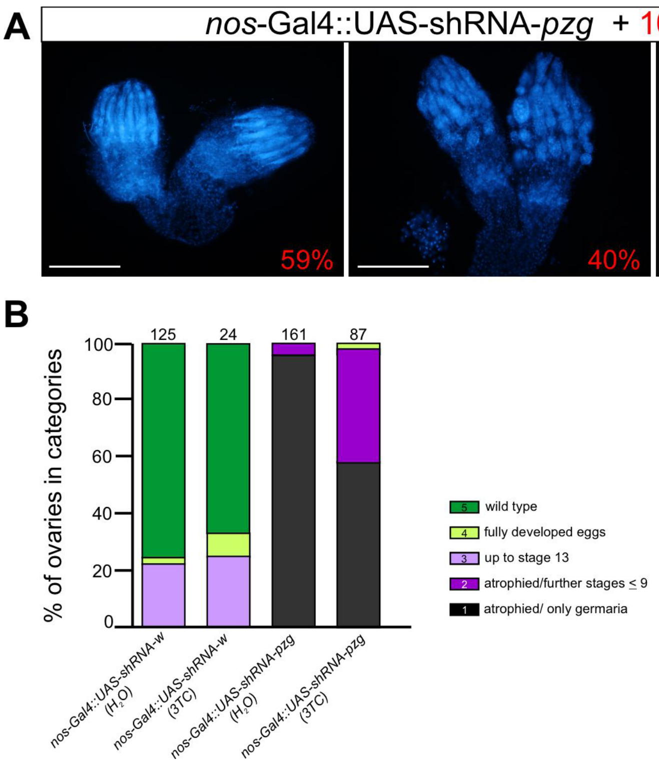


**Figure S1**: *Inhibition of reverse transcriptase ameliorates the pzg mutant phenotype* Feeding of Lamivudine (3TC), an inhibitor of reverse transcriptase, during larval development improved the atrophied phenotype of *nos*-Gal4::UAS-shRNA-*pzg* ovaries. (**A**) DAPI-stained examples for atrophied germaria (left), development of further stages up to stage 9 (about 40% of the females; middle), and germaria with occasional fully developed eggs (about 1%; right). Scale bars, 200 µm. (**B**) Statistics of 3TC feeding assay with respective controls. Number of analyzed ovaries is given above each bar; the respective genotype is indicated below. Categories are: **1 (black)**: atrophied ovaries, i.e. only structures resembling germaria; **2 (dark purple)**: rudimentary ovaries with some follicles developed up to stage 9; **3 (light purple)**: development up to stage 13 in some ovarioles; **4 (light green)**: complete egg development can be observed; **5 (dark green)**: like a wild type ovariole.

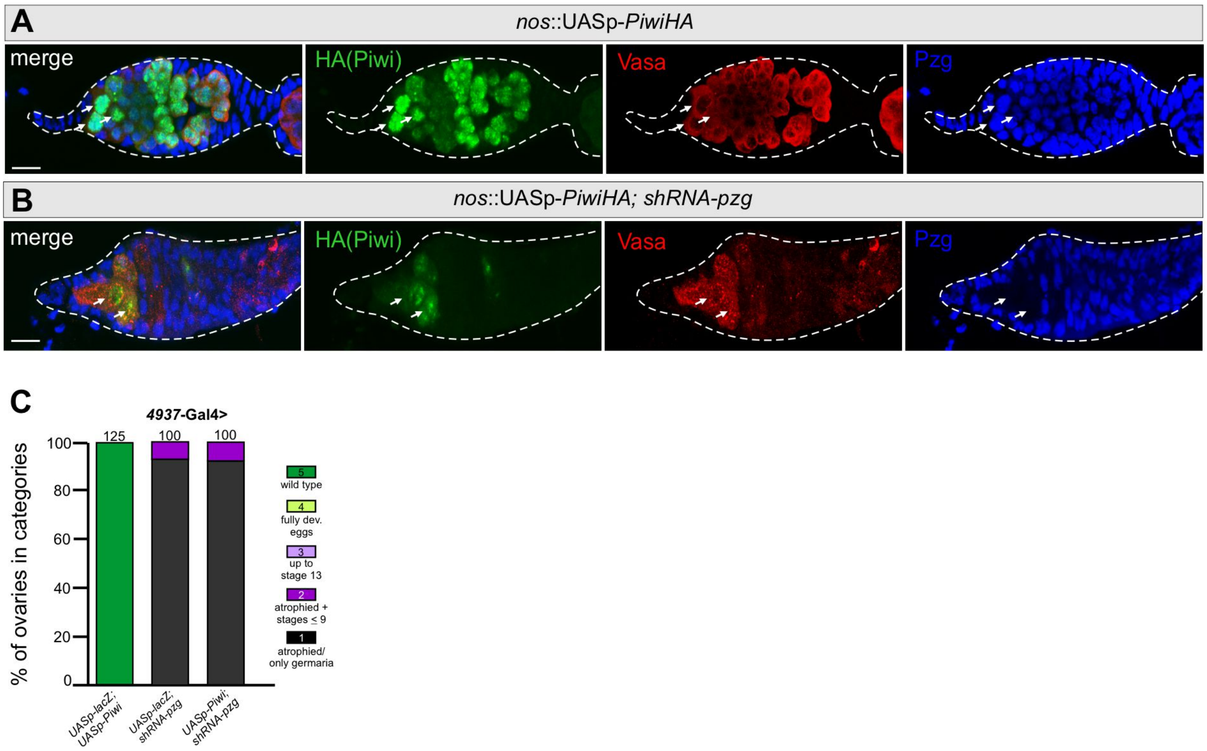


**Figure S2**: *Rescue assays with Piwi overexpression*

(**A**) Ectopic expression of Piwi in germline cells (*nos*-Gal4::UASp-*piwi-HA)* results in Piwi accumulation in GSCs (green, arrows). Germ cells labelled with Vasa (red); nuclei labelled with Pzg (blue). (**B**) Upon Pzg depletion (*nos*-Gal4::UASp-*piwi-HA; UAS*-shRNA-*pzg*), ectopic Piwi protein is barely detected in GSC nuclei (arrows). Co-staining of HA-tagged Piwi (green), Vasa (red) and Pzg (blue) in 0-3 days old germaria using the respective antibodies. Scale bars, 10 µm. (**C**) Rescue assays of *pzg* atrophic ovary phenotype by overexpressed Piwi in the germline. Ovaries derived from 3-5 days old females of the indicated genotype were classified in 5 categories according to their state of development. Number of analyzed ovaries is given above the bars; the examined genotype is given below.

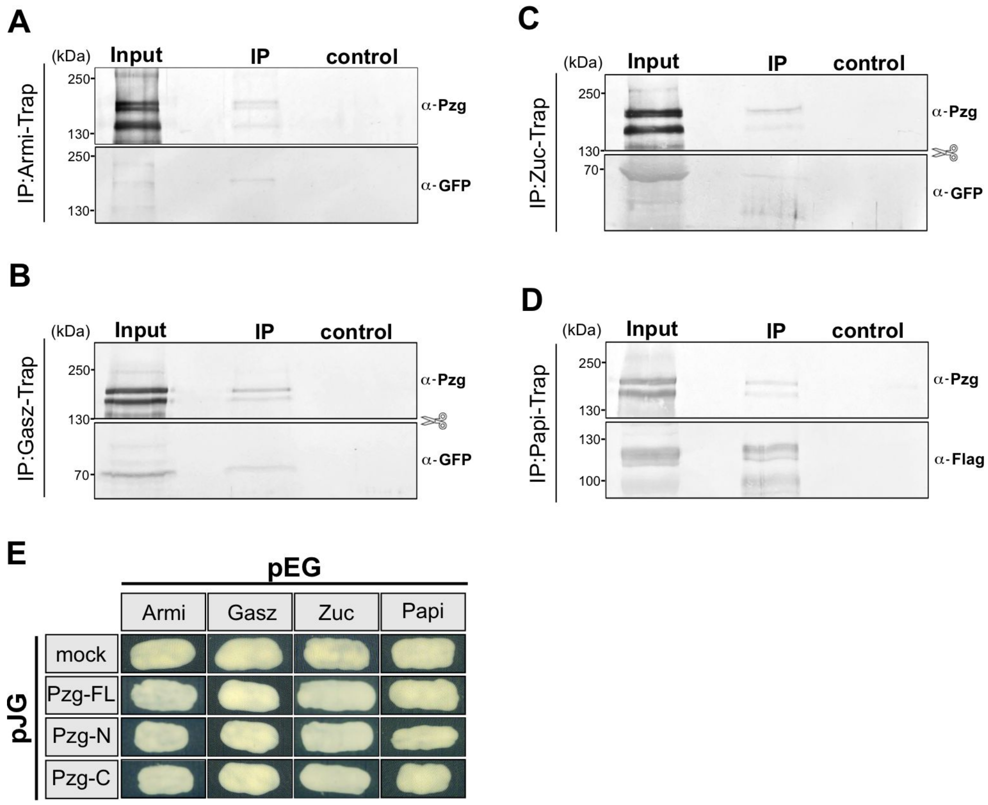


**Figure S3**: *Pzg's association with factors of the piRNA processing machinery*

(**A-D**) Western Blot analysis of co-immunoprecipitations using tagged protein, GFP-Armi (A), GFP-Gasz (B), GFP-Zuc (C), or Flag-Papi (D) from ovarian extracts. Pzg was detected with specific antibodies. 10% of the Input used for the IP-traps were loaded for comparison; control describes the Trap with agarose beads only. (**E**) Yeast two-hybrid interaction assays revealed no direct protein-protein interactions between Pzg and Armi, Gasz, Zuc or Papi.

**Figure S4**: *Uncropped blots of co-immunoprecipitations*

*Uncropped blots from Fig. 6A:* Blots were sliced before antibody treatment in Rhi-, Moon-, and Trf2-trap experiments, indicated by a schematic scissor in Fig. 6A.

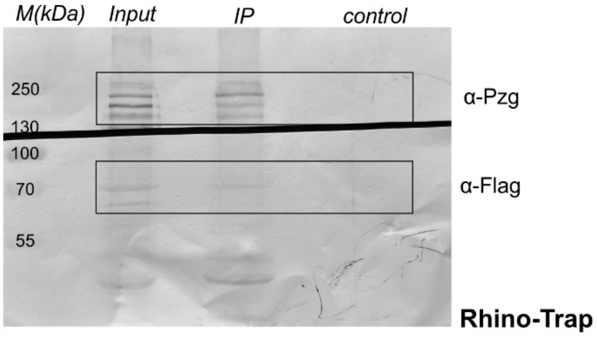


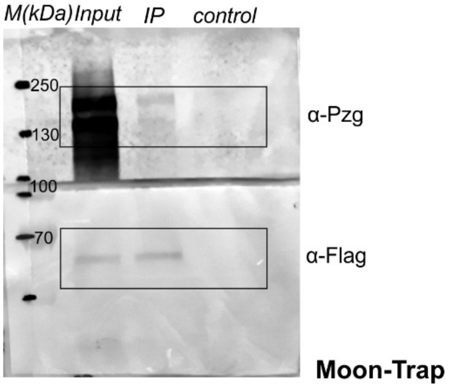


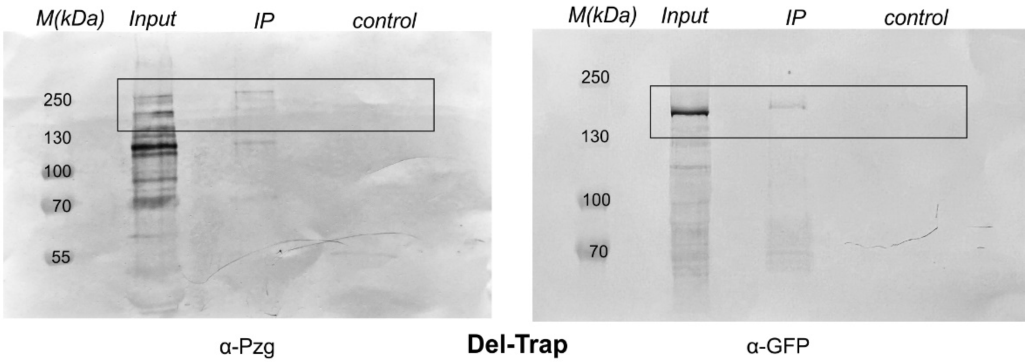


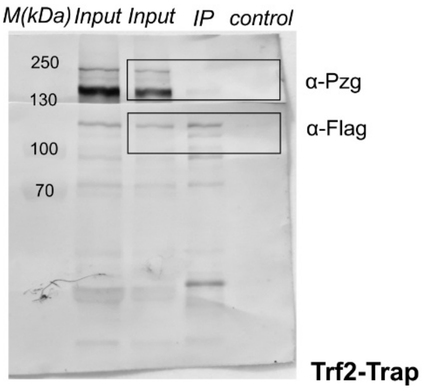

*Uncropped blots from Fig. 8C,F:* Blot was sliced before antibody treatment in Piwi, indicated by a schematic scissor in Fig. 8C.

**C**

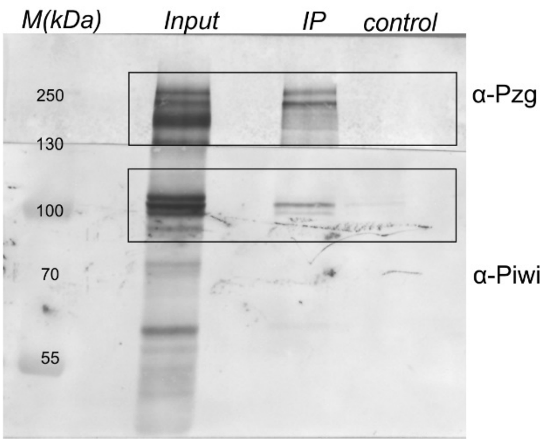


**F**

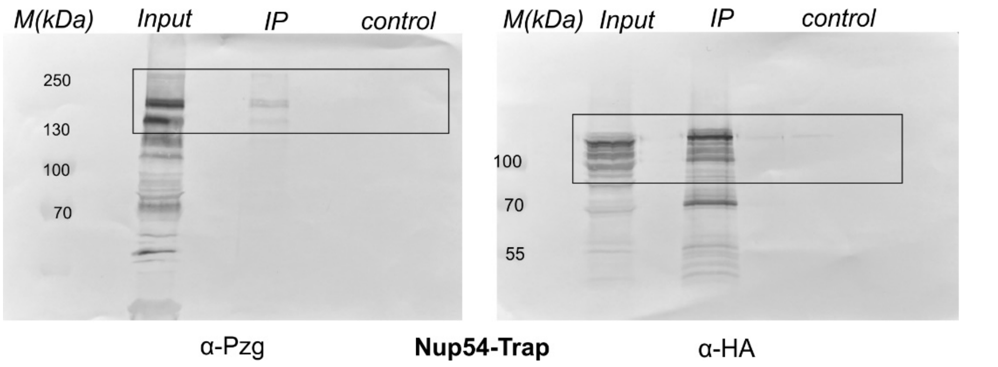


*Uncropped blots from Fig. 9A,B:* Blot was sliced before antibody treatment in Vasa, indicated by a schematic scissor in Fig. 9B.

**A**

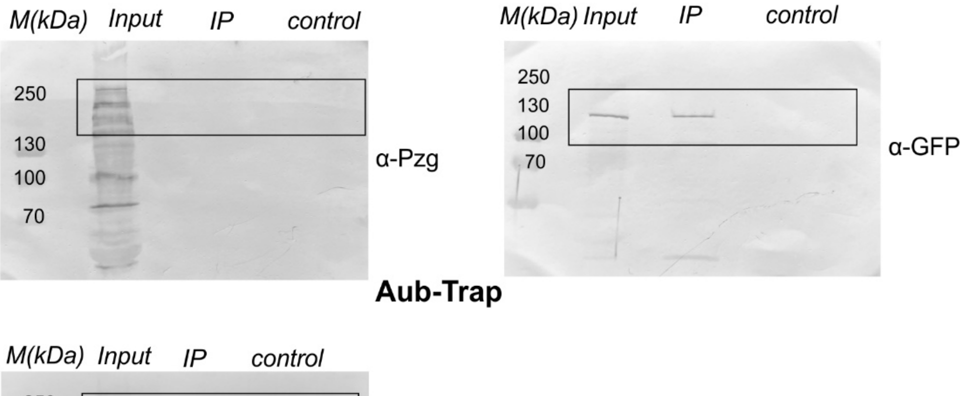


**B**

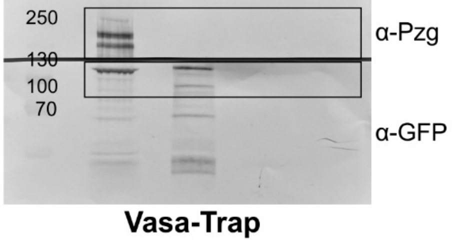

*Uncropped blots from Fig. 10A-D:* Blots were sliced before antibody treatment in Mael-, Pzg-, and HP1a-trap- experiments, indicated by a schematic scissor in Fig. 10A,C,D.

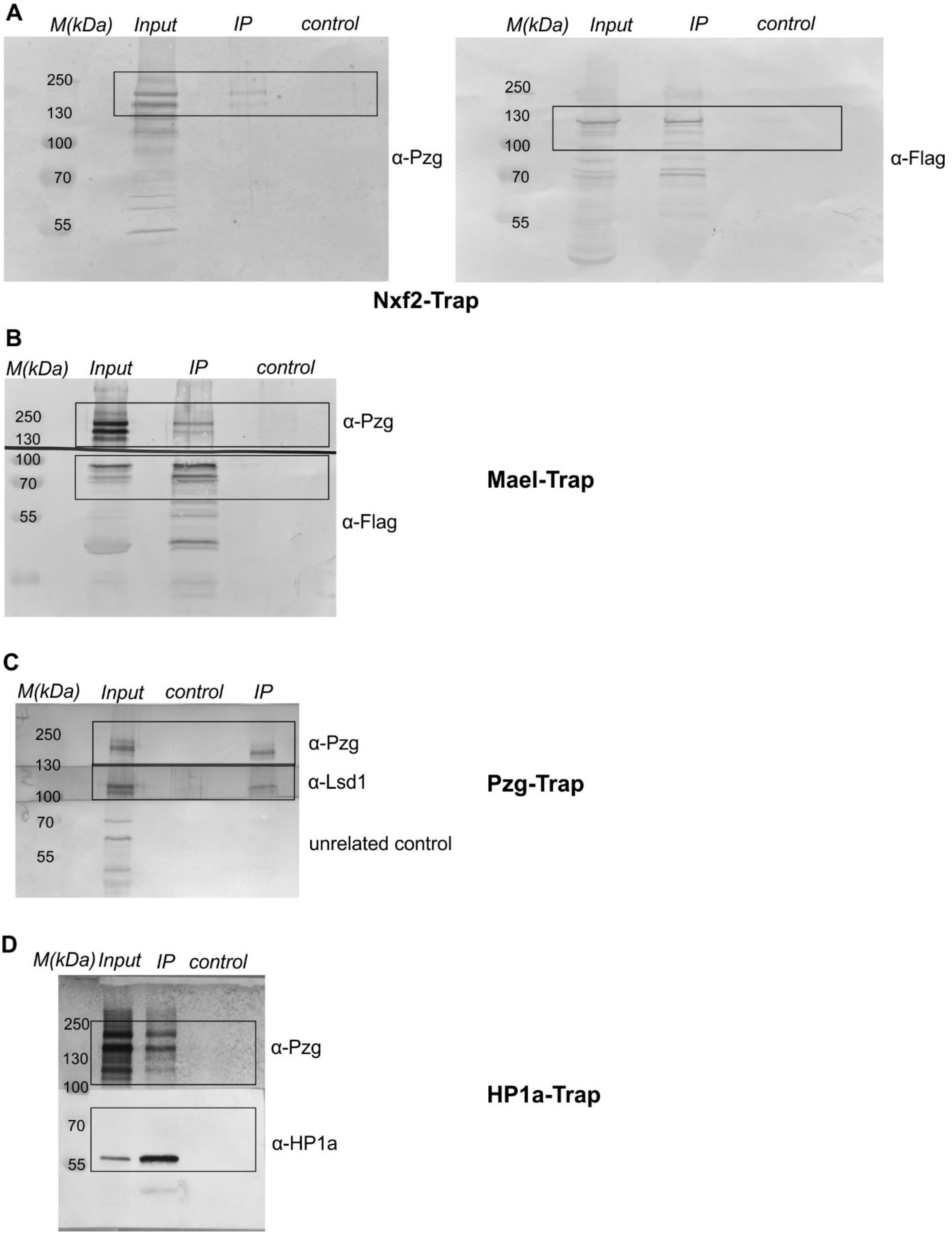

*Uncropped blots from Fig. S3* Blots were sliced before antibody treatment in Gasz-trap and Zuc-trap experiments, indicated by a schematic scissor in Fig. 9S1B,C.

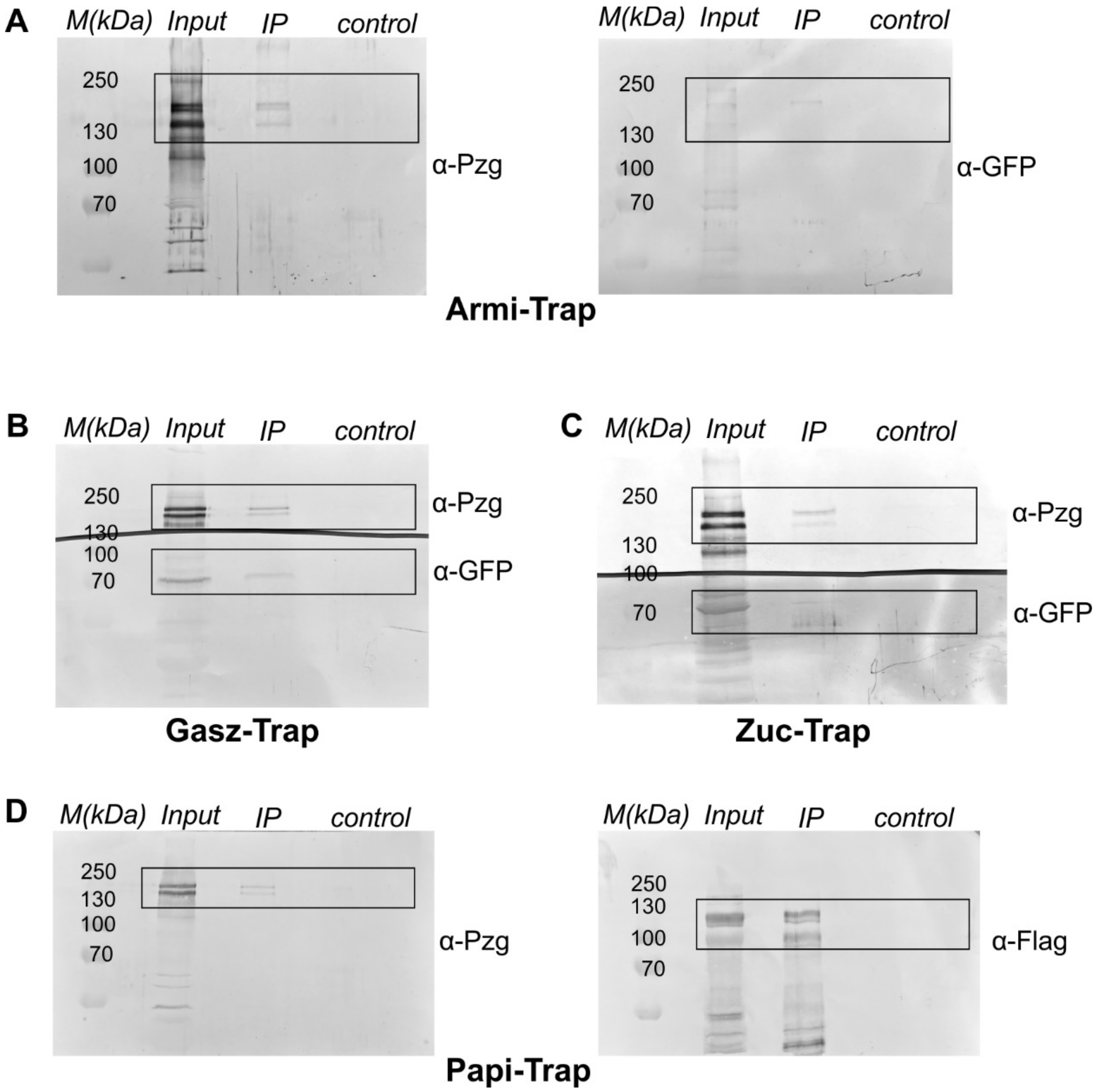
